# Plasmid partitioning driven by collective migration of ParA between nucleoid lobes

**DOI:** 10.1101/2023.10.16.562490

**Authors:** Robin Köhler, Seán M. Murray

## Abstract

The ParABS system is crucial for the faithful segregation and inheritance of many bacterial chromosomes and low-copy number plasmids. However, despite extensive research, the spatio-temporal dynamics of the ATPase ParA and its connection to the dynamics and positioning of the ParB-coated cargo has remained unclear. In this study, we utilise high-throughput imaging, quantitative data analysis, and computational modelling to explore the *in vivo* dynamics of ParA and its interaction with ParB-coated plasmids and the nucleoid. As previously observed, we find that F-plasmid ParA undergoes collective migrations (‘flips’) between cell halves multiple times per cell cycle. We reveal that a constricting nucleoid is required for these migrations and that they are triggered by a plasmid crossing into the cell half with greater ParA. Using simulations, we show that these dynamics can be explained by the combination of nucleoid constriction and cooperative ParA binding to the DNA, in line with the behaviour of other ParA proteins. We further show that these ParA flips act to equally partition plasmids between the two lobes of the constricted nucleoid and are therefore important for plasmid stability, especially in fast growth conditions for which the nucleoid constricts early in the cell cycle. Overall our work identifies a second mode of action of the ParABS system and deepens our understanding of how this important segregation system functions.

## Introduction

In bacterial cell division, the accurate segregation and positioning of chromosomes and plasmids is essential for their faithful inheritance. Indeed, with only a few copies per cell, random positioning cannot ensure the stability of low copy-number plasmids and an active partitioning system is required. For this, many plasmids and bacterial chromosomes use the widespread ParABS partitioning system (1, 2), which is composed of three components: (i) *parS*, a centromere-like binding site, (ii) ParA, a deviant Walker-type ATPase, and (iii) ParB, a protein that binds to the *parS* site. ParA, in its dimeric ATP-bound state, binds DNA non-specifically, coating the nucleoid of the cell (3, 4). It detaches from the DNA upon hydrolysis and requires a prolonged cytosolic conformational transition before again becoming competent for binding (5, 6). ParB, which also dimerizes, binds to *parS* and spreads out along the DNA for several kilobases to form the nucleoprotein ‘partition complex’, visible using fluorescent microscopy (7, 8). Very recently, it was found that the ParB of several species/plasmids is a CTPase that can entrap and slide along the DNA, potentially explaining the observed spreading (9–16). ParB also stimulates the ATPase activity of ParA, generating a local depletion of ParA-ATP around the plasmid (17–19). The resulting ParA-ATP gradient, shaped by both the nucleoid and the plasmids is then believed to direct plasmid movement and result in the regular (equi-distant) positioning of plasmids across the length of the cell (6, 17, 18, 20).

At the molecular level, the central hypothesis for how directed movement occurs is the existence of protein tethers between ParB bound to the cargo (the *parS*-proximal DNA of the plasmid or chromosome) and non-specifically bound ParA-ATP on the underlying nucleoid. The creation of these tethers, which are transient due to ParB stimulated hydrolysis, depends on the local concentration of nucleoid-bound ParA such that more tethers are created in the direction of increasing ParA concentration, thereby providing the means for the cargo to sense the gradient (21, 22). The pulling force itself is believed to be generated by the elastic fluctuations of the DNA and/or the protein tethers as suggested by computational modelling (23–26).

However, while *in vitro* reconstitutions support this view (19, 27, 28), the nature of the plasmid movement *in vivo* and its connection to ParA are much less understood. First, regarding the plasmid, while both oscillatory and defined positioning have been described, the prevalence of either mode within and across species was unclear (17, 18, 29, 30). In this direction, we recently used high-throughput imaging and analysis to quantify the dynamics of F-plasmid and found that it is elastically ‘pulled’ to defined locations within the cell (mid-cell for a single plasmid, the quarter positions for two plasmid) (31). However, at the lowest plasmid concentrations (a single plasmid in a long cell) the dynamics become oscillatory in nature. This was even more pronounced for the plasmid pB171 harbouring a distantly related ParABS system, which exhibits clear oscillatory dynamics across the length of the cell in almost all cells containing a single plasmid (31).

Several models that can explain (typically a subset of) the observed plasmid dynamics have been proposed, both separately from (21, 22, 32, 33) and including (23, 26), the molecular mechanism of force generation. We combined aspects of these models into a new stochastic model that could produce all the dynamical regimes within a single unifying model (31). Critical for this was the understanding of the importance of the length-scale of ParA diffusion on the nucleoid as the primary parameter controlling the transition between oscillatory dynamics and targeted positioning. However, while a quantitative comparison of this model with measurements of F plasmid gave remarkable agreement, a similar comparison to the ParA distribution was not performed.

Indeed, the dynamics of ParA and its correlation to plasmid movement has remained one of the least understood aspects of this system. While plasmids have been observed to follow a retracting gradient of ParA across the cell (17, 18, 20) similar to an *in vitro* reconstitution (19), the connection between the two components in individual cells is often unclear. In particular, the ParA of F plasmid, TP228 and most recently ParA2 of *Vibrio cholerae* have been observed to flip between the two halves of the cell (5, 34, 35). However, none of the current models, including our own, can reproduce this behaviour, indicating a fundamental gap in our understanding of how this system functions.

Here, we investigate ParA dynamics and its connection to F plasmid movement using a combination of high-throughput imaging and quantitative data analysis. We find that flips of ParA between the two cell halves occur only after the nucleoid has begun to constrict and appear to be triggered by the movement of a plasmid from the cell half with the least ParA to the cell half with the most. This is most likely to occur when there is an asymmetry in the amount of plasmids/ParB in each cell half and the effect of the flips is to encourage equi-partitioning between the cell halves. Building on our previous stochastic model of plasmid positioning, we show that this behaviour can be explained by cooperative ParA binding to the nucleoid as has already been suggested for other ParA proteins (5, 35–37), and for membrane recruitment of the ParA-like protein MinD (38, 39), which also exhibits oscillatory dynamics. Using our model, we demonstrate that ParA flipping reduces plasmid loss in conditions with early nucleoid constriction by ensuring partitioning of plasmids between the two lobes of the nucleoid. This is especially important for cells born with a single plasmid in which the nucleoid sufficiently constricts before the plasmid replicates. Thus the ParABS system has two modes of action to effect plasmid stability: 1) regular positioning of plasmids on (each) nucleoid mass and 2) equi-partitioning of plasmids between the two lobes of the constricted nucleoid. These findings advance our knowledge of plasmid partitioning, while also being of relevance for our understanding of chromosomal ParABS systems.

## Results

### ParA flips abruptly between cell halves

To study the dynamics of F plasmid ParA and its relationship to plasmid positioning and the cell cycle, we used our previously developed high-throughput microfluidic and image analysis pipeline (31) (Fig. S1). We visualised the two protein components using an F plasmid derivative carrying functional fluorescent fusions of ParA and ParB (ParA-mVenus, ParB-mTurquoise2) (40). Given that ParB foci separate within 5 minutes of plasmid replication (41, 42), we assume in the following that each focus represents a single plasmid (43). The nucleoid of the host *E. coli* cell was also imaged using chromosomally expressed HU-mCherry (44). A representative cell cycle is shown in Figure 1A. We found that while ParA is distributed relatively uniformly at birth, later (at ∼30 min in the example shown) it often undergoes collective migrations between the two halves of the cell, between which it remains in a stable asymmetric state with the majority of ParA in one cell half. These observations are consistent with previous reports (17, 34). We term these migrations ‘flips’ to distinguish them from wave-like oscillations, which are also observed in ParABS systems, most notably that of the plasmid pB171 (18, 21).

**Fig. 1.**
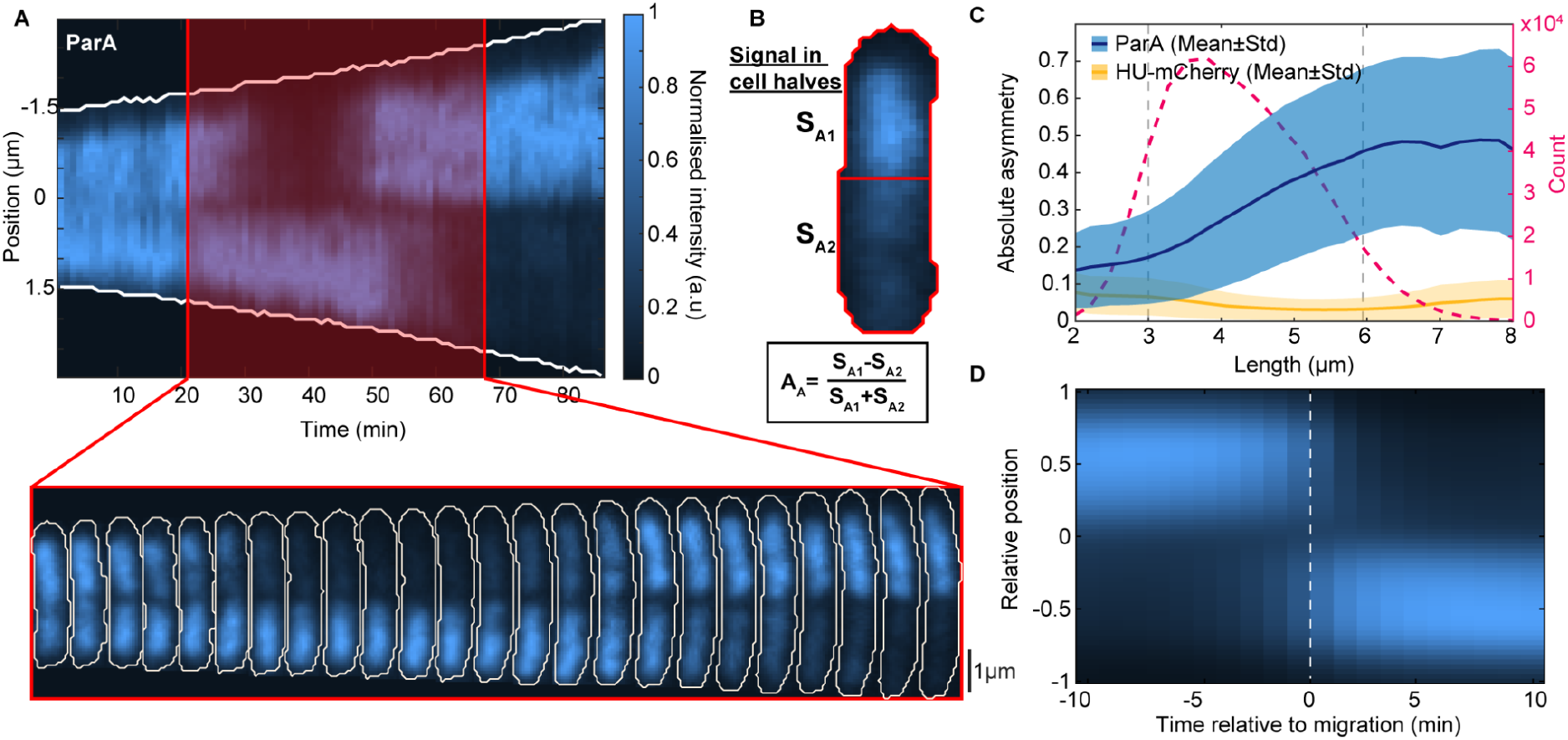
Characterising ParA flips. **(A)** Top: Kymograph of ParA-mVenus along the long axis of a cell during its cell cycle (1 minute time intervals). Bottom: ParA-mVenus signal within the cell for every second frame of the red highlighted region. **(B)** The ParA-mVenus signal in each half (*S*_*A*1_, *S*_*A*2_) is used to calculate the ParA asymmetry: 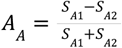. A value of 0 corresponds to an even distribution, +1/-1 correspond to all of the signal coming from one or the other cell half. **(C)** Blue: Absolute ParA asymmetry of cells plotted against length (mean ± SD). Orange: Absolute asymmetry of the nucleoid (HU-mCherry) measured as for ParA is shown for comparison. Red: Number of data points at each length. **(D)** Average kymograph of all ParA flips aligned by the moment the flip occurred. Length was normalised such that +1/-1 correspond to the poles of the cell. Furthermore, the cells were orientated such that the majority of ParA is in the same cell half before the flip. Panel **(C)** was generated from 5219 cell cycles and panel **(D)** was generated from 2537 flipping events.

To quantify ParA asymmetry, we first measured the fractional difference *A*_*A*_ in the ParA-mVenus signal between each half of the cell (Fig. 1B). We found this to increase significantly with cell length, particularly between the average birth and division lengths of 3 μm and 6 μm respectively (Fig.1C, S1D) before plateauing at a value (∼0.5) corresponding to an average 75:25 distribution of ParA between cell halves. By defining ParA to be in an asymmetric state from at least a 67:33 distribution (|*A*_*A*_| > 0. 33), we could also identify flips by the switching of ParA between asymmetric states (Fig. S2A). According to this definition, cells typically display two flips during their cell cycle (Fig. S2B), occurring, on average, 20 minutes apart but with substantial variation (Fig. S2C). The time between these flips lengthens as the cell grows (Fig. 2SD), suggesting that the asymmetric state becomes increasingly stable (less likely to flip) as the cell grows. ParA asymmetry at division often results in one daughter cell inheriting the majority of ParA, producing a very heterogeneous distribution of in newborn cells: on average one daughter cell inherits 73% of the ParA signal (Fig. S2E). However, while the ParA signal is certainly noisier in cells with little ParA, flips occur across the entire range of ParA levels (Fig. S3A). Indeed, beyond a point, ParA asymmetry appears to be independent of ParA levels since the mean absolute ParA asymmetry |*A*_*A*_| is stable across the upper half of the distribution (Fig. S3C).

**Fig. 2.**
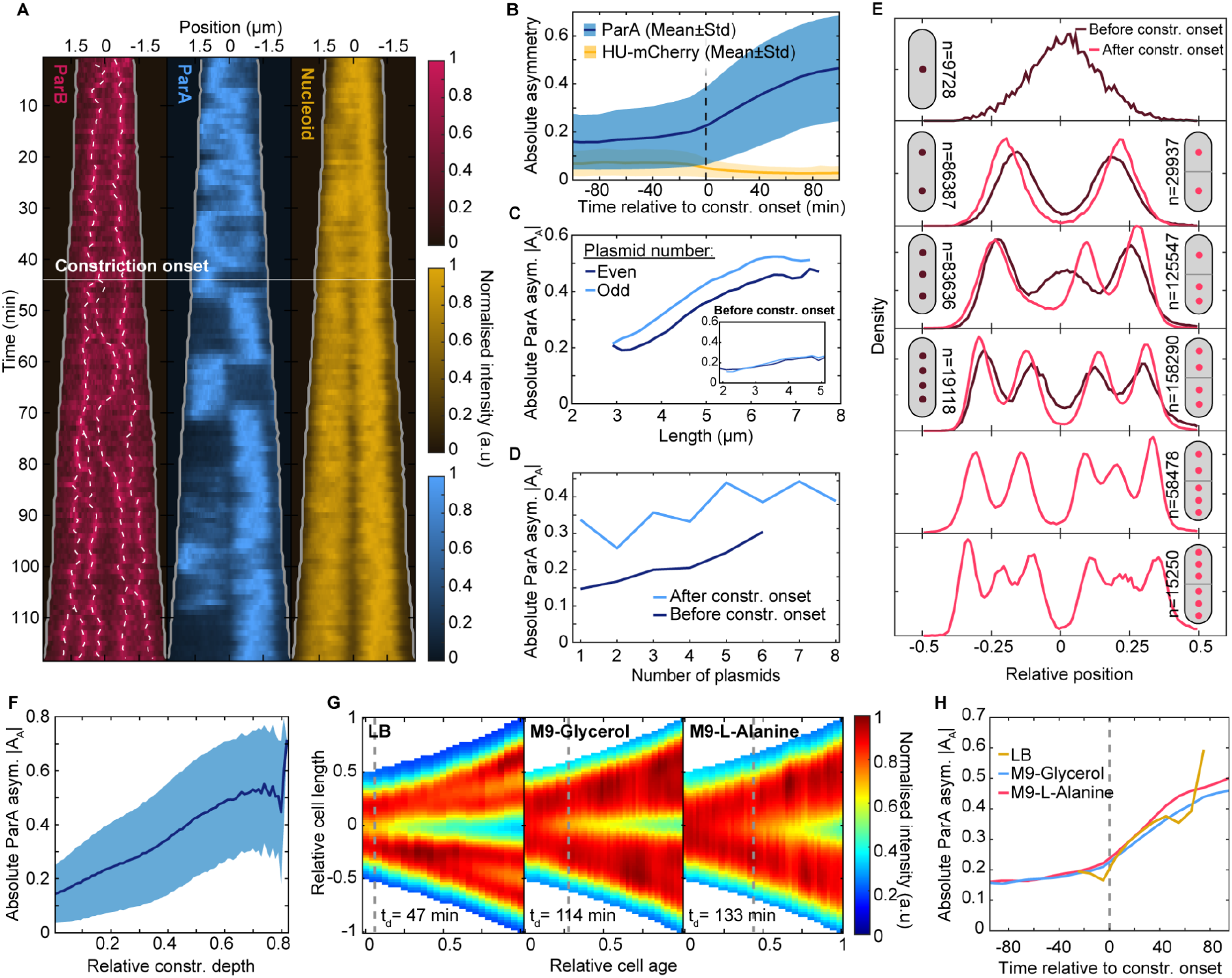
ParA asymmetry increases from the onset of nucleoid constriction. **(A)** Kymograph of Nucleoid (HU-mCherry), ParA-mVenus, ParB-mTurquoise2 signals along the long cell axis during an example cell cycle (1 minute time intervals). White dashed lines indicate tracked foci. The solid white line indicates the detected onset of nucleoid constriction. **(B)** Average absolute ParA (blue) and nucleoid (orange) asymmetry relative to the onset of nucleoid constriction (mean ± std). Nucleoid asymmetry was calculated similarly to ParA asymmetry in Figure 1B. **(C)** The absolute asymmetry of ParA in cells containing varying numbers of plasmids after and before (inset) the onset of nucleoid constriction. **(D)** The absolute asymmetry of ParA in cells with different numbers of plasmids after the onset of the nucleoid constriction. **(E)** Histograms of the distribution of plasmids before (dark red) and after (red) constriction onset. Cells after constriction onset are aligned such that the side with the most plasmids is on the right. The number next to each cartoon shows the number of data points for each configuration. **(F)** Absolute ParA asymmetry (mean ± std) plotted against relative constriction depth. **(G)** Averaged kymographs of the nucleoid (HU-mCherry) in three different growth conditions. **(H)** Mean absolute ParA asymmetry as in panel **(B)** for all three growth conditions studied. The data in panels **(A)**-**(F)** is from the M9-Glycerol condition as in Figure 1. Number of cell cycles used: 5219 (M9-Glycerol), 606 (LB) and 569 (M9-L-Alanine).

The example cell cycle in Figure 1A, also highlights another feature of ParA dynamics. During a flipping event, the ParA signal often appeared to decrease uniformly in one half, while simultaneously increasing uniformly in the other. This was corroborated by averaging over many flipping events (Fig. 1D) and by imaging cells on an agarose pad at higher time resolutions (Fig. S4). The migration occurs rapidly, taking only a few minutes. While we also observed cases of oscillatory, retracting-gradient-like ParA dynamics (Fig. S5A), similar to that described for the pB171 (18), these cases were rare, occurring in only approximately 3% of cells.

It is important to note that ParA could also exhibit heterogeneity early in the cell cycle (Fig. S5B), as quantified by comparison to the asymmetry of the largely symmetrical nucleoid signal (Fig. 1C). However, this tended to be weaker, more heterogeneous and transient and without the step-like profile and mid-cell demarcation that occurs later in the cell cycle. Similar heterogeneity could also sometimes be observed within the lobes of a constricted nucleoid (Fig. S5B, Fig. 2A). Thus the asymmetry and flips of ParA between cell halves that we study here is in addition to, and not a complete description of, the inherent heterogeneity of the system.

### ParA asymmetry increases upon nucleoid constriction

We next investigated what underlies the increase of ParA asymmetry with cell length (Fig. 1C). Since the flipped ParA state is demarcated by mid-cell, we wondered if the constriction of the nucleoid may play a role. Indeed, we found ParA flips to be stronger and more frequent after the nucleoid begins to constrict (Fig. 2A, S3A). To quantify this, we first set a threshold depth to characterise dips in the HU-mCherry signal (Fig. S6). Since the nucleoid is dynamic and shallow constrictions can be transient, we then determined, for each cell cycle, the onset of nucleoid constriction i.e. the time point from which a constriction deeper than the threshold is continuously present for the remainder of the cell cycle. We found that ParA asymmetry remains relatively constant before nucleoid constriction begins but increases abruptly afterwards (Fig. 2B). We found that this transition is not directly due to cell length or plasmid copy number increases since ParA asymmetry is greater after the onset of constriction than before it at the same cell length (Fig. 2C) or plasmid copy number (Fig. 2D). On the other hand, we observed an almost linear relationship between ParA asymmetry and the depth of the nucleoid constriction (Fig. 2F).

If the onset of nucleoid constriction is indeed a prerequisite for ParA flipping, then the relationship to ParA asymmetry should be maintained even when the nucleoid constricts earlier or later in the cell cycle (45). We therefore repeated our measurements with cells grown on a poorer carbon source (M9-L-alanine) and in richer media (LB). In the latter case, nucleoid constriction begins on average shortly after birth or already in the previous cell cycle, whereas using L-alanine constriction begins about halfway through the cell cycle. Our default M9-Glycerol condition lies somewhere in between (Fig. 2G). We found that in both cases ParA asymmetry begins to increase noticeably upon the onset of nucleoid constriction (Fig. 2H). This is despite the large variation in the timing of constriction onset both within and between the populations (Fig. S6F). The curves of all three conditions also collapsed onto each other. We speculate this is due to similar progression of the replication forks and, correspondingly, the progression of nucleoid constriction and hence ParA asymmetry increases similarly across the three conditions.

As a further test, we imaged the nucleoid of cells upon treatment with the antibiotics rifampicin and chloramphenicol and upon thymine depletion (the strain is a thymine auxotroph), all of which are known to affect nucleoid organisation (46, 47). For all three cases, we found that most nucleoid constrictions (relative depth greater than 20% as in Fig. S6) were lost by the end of imaging i.e. nucleoids remerged (Fig. S7A). This was concomitant with a drop in ParA asymmetry, which followed the same linear relationship to constriction depth that we found earlier (Fig. S7B). In individual cells, the loss of ParA asymmetry and flipping upon nucleoid remerging could be clearly seen (Fig. S7C-E). We will show below that a similar behaviour could also be seen in some unperturbed cells. Our data therefore suggest that nucleoid constriction is a requirement for the asymmetry of ParA between cell halves and its associated flipping.

Since plasmid-associated ParB stimulates hydrolysis and DNA unbinding of ParA, unequal plasmid partitioning should also contribute to ParA asymmetry. Indeed, constricted cells with unequally partitioned plasmids have on average greater ParA asymmetry than those with equally partitioned plasmids (Fig. S6G). However ParA asymmetry before constriction is independent of partitioning (Fig. S6G), further indicating the importance of nucleoid constriction. Consistent with these results, after constriction onset ParA asymmetry is slightly but consistently higher in cells with an odd number of plasmids (Fig. 2C,D). Since plasmids are associated to higher density regions of the nucleoid (44), they are naturally biased away from the nucleoid constriction site. Therefore, after constriction cells with odd copy numbers necessarily have unequally positioned plasmids (Fig. 2E, cell with three plasmids).

### Plasmid-bound ParB triggers ParA migration

We noticed that in many cell cycles ParA flipping was coincident with a plasmid moving from one half of the cell to the other (Fig. 3A, also Fig. 2A, S2A, S3, S5). To quantify this, we analysed plasmid movement and replication in the ten minutes preceding each flip (Fig. 3B-F). We found that in 63% of cases, a change in the plasmid distribution between the cell halves could be observed, either due to a plasmid crossing mid-cell or due to plasmid replication. For the remaining 37%, no change in the plasmid distribution was detected i.e. ParA could be asymmetric and flip even when plasmids were equally partitioned (Fig. 3E).

**Fig. 3.**
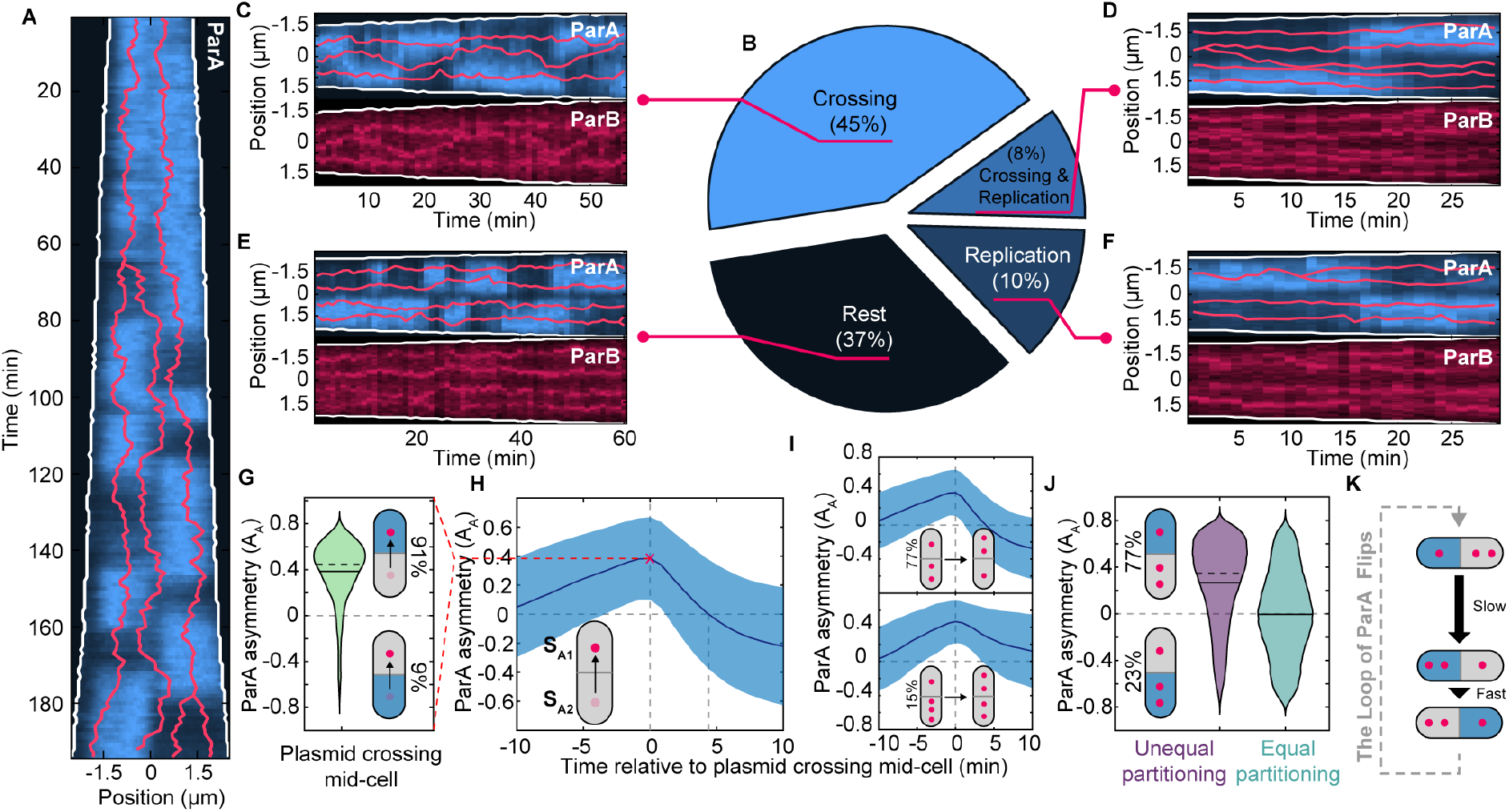
Mid-cell plasmid crossing precedes ParA flips. **(A)** Kymograph of ParA-mVenus. Red lines indicated tracked ParB-foci. **(B)** Pie chart of which cellular event precedes (within 10 min) a ParA flip: a plasmid crossing mid-cell (Crossing), plasmid replication (Replication) or neither crossing nor replication (Rest). **(C)-(F)** An example portion of a cell cycle in which the indicated event preceded ParA migration. **(G)** Distribution of ParA asymmetry at the time of a plasmid crossing event. **(H)** ParA asymmetry (mean ± std) in the 20 minutes surrounding a plasmid crossing, with 0 marking the time a plasmid crosses mid-cell. Positive asymmetry corresponds to more ParA in the cell half the plasmid crosses into. **(I)** Same as **(H)**, but data were binned based on the plasmid configuration before the crossing. The top panel includes crossings that do not change the disparity in plasmid number (e.g. from 2:1 to 1:2, or 3:2 to 2:3). The bottom panel includes crossing events that equalise the number of plasmids in both cell halves (e.g. from 3:1 to 2:2, or 4:2 to 3:3). **(J)** ParA asymmetry is binned by equal and unequal plasmid partitioning. For unequal partitioning, positive asymmetry values correspond to more ParA in the cell half with less plasmids as indicated. **(K)** Proposed order of events. **(G)**-**(J)** considers only crossing events after the onset of nucleoid constriction. **(B)** and **(G)**-**(J)** were generated using 5219 cell cycles grown on M9-Glycerol. Solid and dashed lines in **(G)** and **(J)** indicate mean and median respectively.

Since our identification of flips requires an arbitrary threshold on the ParA asymmetry *A*_*A*_, we switched to focusing on plasmid midcell-crossing events, which can be identified more easily. In 91% of cases, the plasmid moves from the cell half with the least ParA to the cell half with the most (Fig. 3G) and the crossing coincides with the time of maximum ParA asymmetry (Fig. 3H). On average, the asymmetry then decreases, reaching parity 5 minutes after the crossing event before increasing in the opposite direction (Fig. 3H). This was predominantly (77%) due to crossings that reversed the asymmetry in the number of plasmids (Fig. 3I), consistent with most (78%) plasmid crossing events being followed by a flip. The average ParA asymmetry decreased more slowly when the number of plasmids in the two cell halves was equalised by the crossing event indicating flips were less likely to occur. Given that the predominant configuration in cells with unequal plasmid partitioning has ParA distributed more to the cell half with fewer plasmids (Fig. 3J), the data indicates an ordering for the plasmid-crossing-induced flipping of ParA (Fig. 3K). Specifically, a plasmid from the cell half with the majority of plasmids moves to the other half of the cell where the ParA concentration is greatest. The resulting relocation of plasmid-bound ParB induces ParA to rapidly flip to the other cell-half reestablishing the initial configuration, presumably through its stimulation of ParA hydrolysis and facilitated by the constriction of the nucleoid. This was supported by an analysis of how the asymmetry of both proteins are related in time, which showed that ParB indeed follows ParA (Fig. S8) and is conceptually similar to the membrane oscillations of the ParA-like protein MinD between the *E. coli* cell poles due to its recruitment of its inhibitor MinE (48).

### Cooperative DNA-binding explains ParA flipping

Based on our results thus far, we suggest the following explanation for ParA flipping. Nucleoid constriction insulates the two lobes from each other by reducing the diffusion of nucleoid-associated ParA dimers between them (28, 49). Thus, in cells with a constricted nucleoid and unequally partitioned plasmids, the different overall ParB-induced hydrolysis rates on each nucleoid lobe leads to the dissociation of ParA from the lobe with more plasmids and its accumulation on the lobe with fewer. Notwithstanding any heterogeneity of the ParA distribution on each lobe, this tends to result in a sharp, almost step-like gradient of ParA across the constriction, or equivalently a strong directed flux of ParA across mid-cell (Fig. S9). We speculate that this flux directs the movement of a midcell-proximal plasmid from the nucleoid lobe with more plasmids and less ParA to the lobe with fewer plasmids and more ParA through, as has been proposed, the formation of ParA-ParB protein tethers connecting the plasmid to the underlying nucleoid (23, 26, 31).

However, to our knowledge, none of the existing models of the ParABS system, including our own, display ParA flips of the type described here but rather only wave-like oscillations. If these occur, they are concomitant with oscillatory movement of (all of) the plasmids and are not demarcated by midcell (26, 31). In our experiments, we see the majority of plasmids in the cell apparently unaffected by the often extreme difference in ParA levels in each cell half with plasmids remaining regularly positioned within each nucleoid. As described above only the movement of a plasmid across the mid-cell appears to correlate with ParA flipping. This indicates that some ingredient is missing from the current models.

The bistable nature of the ParA profile (in particular that ParA can be asymmetric even when plasmids are equally partitioned as in Fig. 3E,F) suggests cooperativity may be involved. Indeed, the ParA of the P1 plasmid, *Thermus thermophilus* and *V. Cholerae* have been shown to bind non-specific DNA cooperatively (36, 50, 51, 5). While oligomerization appears to be involved, it is evidently transient as ParA associates uniformly to and from the DNA (5, 6, 27, 28). We therefore studied the effect of cooperative ParA binding on the system by implementing it into our previous ‘Hopping and relay’ model (31); see SI Appendix and Fig. S10 for details. Note that we do not specify the molecular details of the cooperativity i.e. whether bound ParA recruit unbound ParA directly or indirectly e.g. by stimulating the transition of cytosolic dimers to their DNA-binding-competent state (5, 6).

Independent of the exact mechanism, cooperative ParA binding creates a positive feedback loop in which more ParA is recruited to the location where already bound ParA is greatest, namely furthest from any plasmids (Fig. 4A,B). In our simulations, this indeed produced an asymmetric ParA distribution and flip-like migrations (Fig. 4D, ii). However, the asymmetry was relatively weak, not clearly demarcated by mid-cell and there was little effect on the plasmids, which remained equally positioned across the cell. In particular the middle plasmid (of three) was maintained at mid-cell in contrast to what we observe experimentally post-constriction (Fig. 2E).

**Fig. 4.**
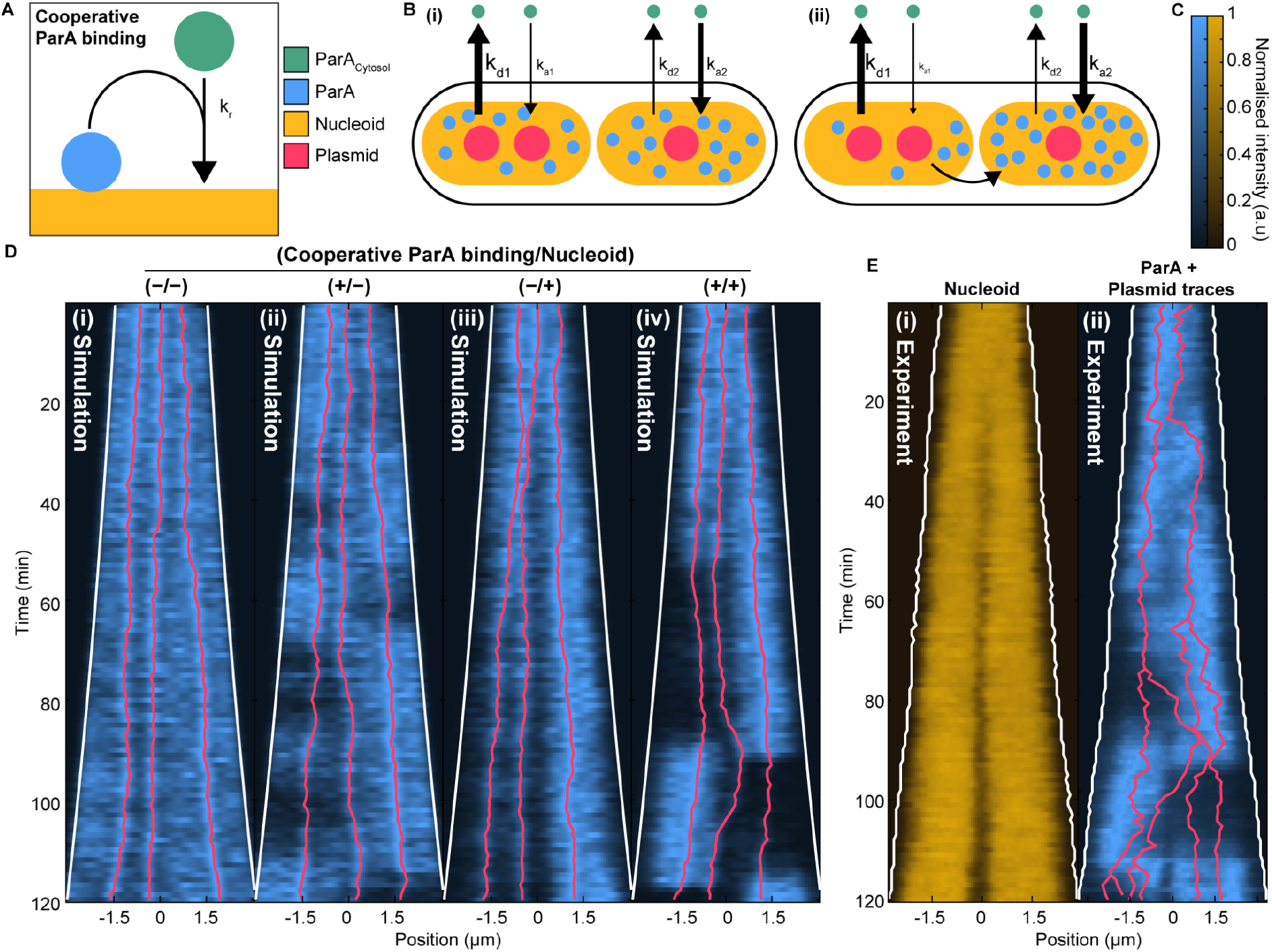
Cooperative ParA binding leads to robust ParA migration. **(A)** A cartoon of nucleoid-bound ParA cooperatively recruiting cytosolic ParA. **(B)** A cartoon of ParA migration in a cell with two nucleoids. **(i)** Plasmid-bound ParB stimulates ParA hydrolysis, increasing the rate of ParA dissociation from the nucleoid. The nucleoid with more plasmids has a higher net dissociation rate and therefore has less ParA bound. **(ii)** The difference in ParA is further increased by cooperative ParA binding. This leads to a sharp gradient or flux of ParA across mid-cell that directs the mid-cell proximal plasmid towards the nucleoid lobe with more ParA. **(C)** Colour scales for **(D) & (E). (D)** Simulated kymographs of ParA performed under four conditions: without any modifications **(i)**, with the incorporation of ParA cooperativity **(ii)**, with the inclusion of a nucleoid model **(iii)**, and with both ParA cooperativity and the nucleoid model **(iv). (E)** Experimental kymographs of the nucleoid **(i)** and ParA **(ii)** signal from an example cell cycle. Plasmid positions are overlaid in red. The nucleoid signal of this cell is used to implement the nucleoid in the simulations shown in **(D, iii & iv)**. See Fig. S10 and methods for details of the parameters used in the simulations.

Since nucleoid constriction is important for both ParA asymmetry and the exclusion of plasmids from mid-cell (as observed post-constriction), we incorporated the nucleoid itself into our simulations using the dynamical 2D HU-mCherry profile from a representative cell (Fig. 4E, i) (see SI Appendix for details). The effect is to bias the diffusion of nucleoid-associated ParA in the direction of greatest nucleoid signal (DNA density), resulting in a decrease of bound ParA at the constriction and of the ability of bound ParA to diffuse across the constriction. The cytosolic pool is unaffected.

First, we examined nucleoid constriction without cooperative binding. We found that after constriction the plasmids separate to each nucleoid lobe with ParA accumulating more on the lobe with fewer plasmids (Fig. 4D, iii). However, this configuration was maintained until the end of the cell cycle and the ParA asymmetry was relatively weak and developed gradually rather than rapidly. Across multiple simulations no ParA flips or plasmid crossing events were observed. On the other hand, when we included both nucleoid constriction and cooperative ParA binding, we found strong ParA asymmetry, the greatest of the three cases, and more realistic plasmid dynamics (Fig. 4D, iv). This meant there was very little ParA in the lower cell half so that the middle plasmid is affected by the flux of ParA coming from the higher cell half and moves stochastically (i.e. could take some time to begin) in that direction. Shortly afterwards, ParA flips to ‘escape’ from the now two plasmids in that cell half and the cycle repeats.

As noted earlier, ParA is also observed to flip even in the absence of any change in the plasmid distribution between the two cell halves (Fig. 3B,E). Such ‘un-induced’ flips mostly occurred when plasmids were equally distributed between cell halves (71% of the cases). To test if our model could reproduce this behaviour we also simulated cells with four plasmids (Fig. S11). Firstly, we found that nucleoid constriction on its own could not generate an asymmetric ParA distribution when the plasmids were equally partitioned, further discounting this scenario. However, in the presence of both nucleoid constriction and cooperative ParA binding, we found that ParA asymmetry and flips could occur even without a change in the plasmid distribution (Fig. S11), consistent with our experimental observations. Thus, while nucleoid constriction is required for the flipped mid-cell-demarcated asymmetric state to exist, cooperativity 1) strengthens the asymmetry which promotes mid-cell crossing and 2) allows this state to exist even when plasmids are evenly partitioned (akin to how stiffness in a see-saw allows it to be stable even when tilted).

To further evaluate our model, we next looked in our data for cell cycles that show unusual nucleoid dynamics that could serve as tests. We identified two such patterns. The first was the transient remerging of two nucleoids (or nucleoid lobes) of the cell (Fig. 5A), which could be due to the action of the DNA repair machinery. Examining ParA in these cells, we found that nucleoid remerging was coincident with the redistribution of ParA to cover the entire remerged nucleoid, before becoming asymmetric again upon nucleoid re-separation. This provides further support for the role of nucleoid constriction in generating ParA asymmetry. When we simulated the plasmid and ParA dynamics of these cell cycles using the measured nucleoid (HU-mCherry) signal and plasmid positions as inputs, we found that we could qualitatively reproduce the same ParA dynamics (Fig. 5B).

**Fig. 5:**
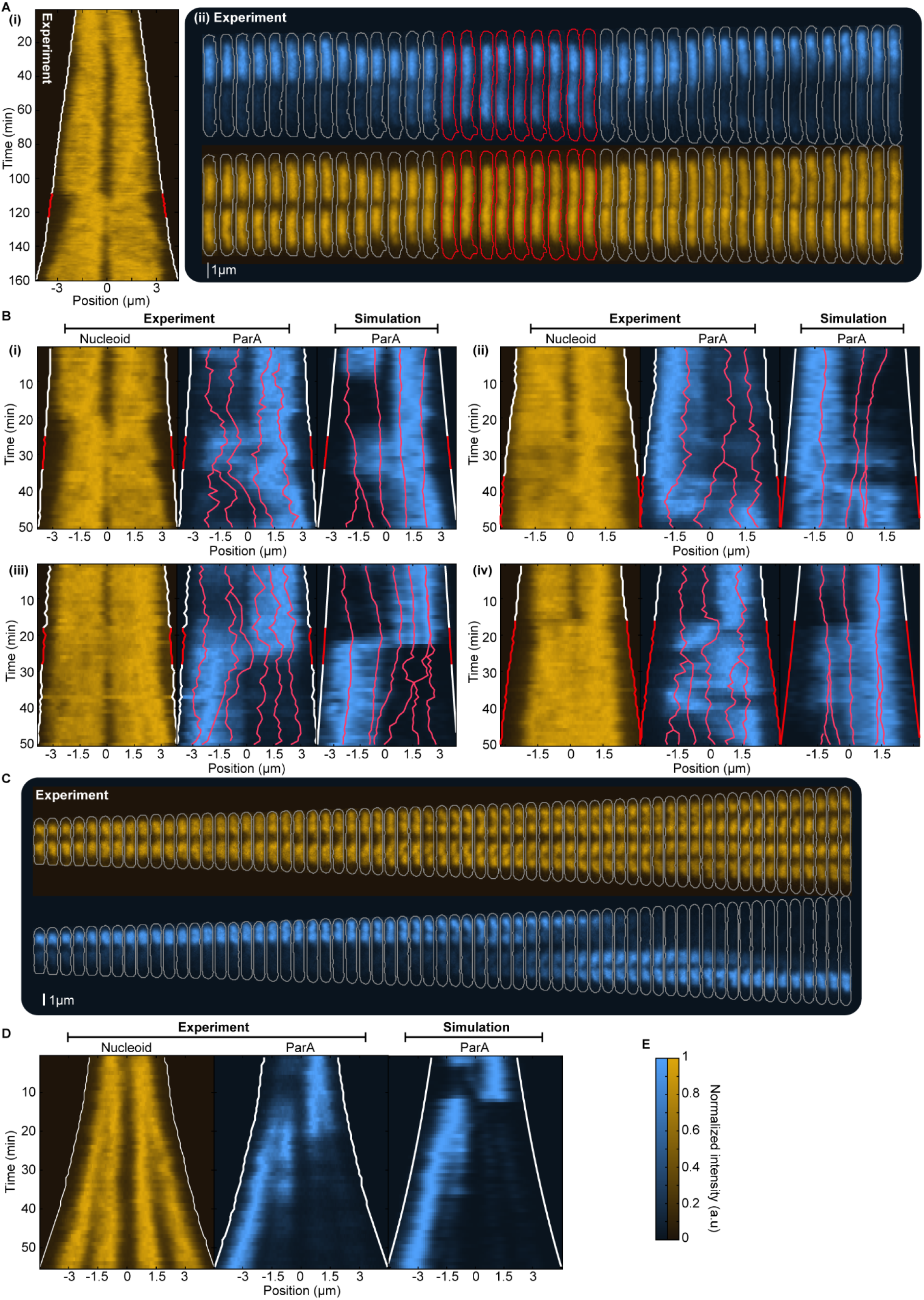
Atypical ParA dynamics are reproduced by our stochastic model. **(A)** Example of transient remerging of the nucleoid. **(i)** Kymograph of the nucleoid signal (HU-mCherry) of a complete cell cycle. White lines indicate the cell border (red when the nucleoid remerges). **(ii)** Fragment of both nucleoid (bottom) and ParA (top) signal of the same cell cycle centred around the remerging event (red cell outlines). **(B)** Four examples of 50 minute fragments of cell cycles in which the nucleoid remerges. Each shows the nucleoid and ParA signal followed by an example simulation using the same nucleoid profile. Plasmid trajectories are displayed as red lines. For **(iii)** 700 ParA dimers were used and 1000 for **(ii) & (iv)**. This was chosen to better match the experimental data. For each of **(i)** to **(iv)**, 300 simulations were performed. **(C)** A complete cell cycle grown in LB, with the nucleoid in orange and ParA in blue. This cell cycle stands out due to its abnormal size and the presence of four clearly separate nucleoids and a strongly localised ParA distribution. **(D)** Kymographs of the nucleoid and ParA signals from **(C)** and a kymograph of simulated ParA using the nucleoid profile from **(C)**. For cells grown on LB media we did not acquire ParB-mTurquoise2 signal; therefore no plasmid trajectories are shown. The plasmid number and initial positions used in the simulations were chosen based on their expected values. **(E)** Colorbars for **(A)**-**(D)**.

The second unusual pattern was observed in cells grown on LB media. In this condition the onset of nucleoid constriction could occur in the previous cell cycle (Fig. 2G) such that cells could have two nucleoid lobes at birth. If those cells in turn divide late, they could have four nucleoid lobes before their division (Fig. 5C). In these cells ParA was very often localised entirely to one of the four nucleoid lobes, typically one of the outer two but could flip between any of them, again indicating the importance of the nucleoid topology. Using the measured nucleoid signal as an input to model, we found that our model could also reproduce this behaviour (Fig. 5D).

Note that, like the real system, our simulations are stochastic and can therefore produce different behaviours for the same input nucleoid profile and initial plasmid positions. However, while there were occasional differences from the *in vivo* observations (e.g. to which nucleoid ParA was localised after re-splitting), the simulations consistently showed good qualitative agreement, supporting our hypothesis that cooperative ParA binding (alongside nucleoid constriction) is a key component for the dynamics of this system.

### The Role of ParA Flips

What is the purpose of the ParA flipping? We hypothesise that it is a mechanism to ensure correct plasmid partitioning after nucleoid constriction has occurred, distinct from the previously described regular positioning of plasmids. Since F plasmids are typically found within the highest density regions of the nucleoid (44), it is likely that they would be excluded from, and have difficulty crossing, the site of nucleoid constriction simply due to the lower amount of DNA and hence ParA there. In richer growth conditions, earlier nucleoid constriction (Fig. 2G, S6F) would make it more likely that all the plasmids are trapped in one nucleoid lobe and therefore destined for the same daughter cell irrespective of their replication. Our results suggest that ParA flipping causes the redistribution of plasmids between the lobes of the constricted nucleoid to ensure accurate inheritance to each daughter cell.

To test if ParA flipping can indeed improve plasmid stability, we incorporated plasmid replication into our stochastic simulations. Focusing first on the extreme case of cells born with a single plasmid, we found that plasmid-crossing-induced flipping of ParA due to cooperative ParA binding improved plasmid partitioning and hence stability (Fig. 6A). Without cooperative binding, plasmids were not always redistributed between the two cell halves, especially when replication occurred after the onset of nucleoid constriction, and ParA was simply maintained in the cell half with fewer plasmids. As a result, plasmid loss occurs (Fig. 6A).

**Fig. 6.**
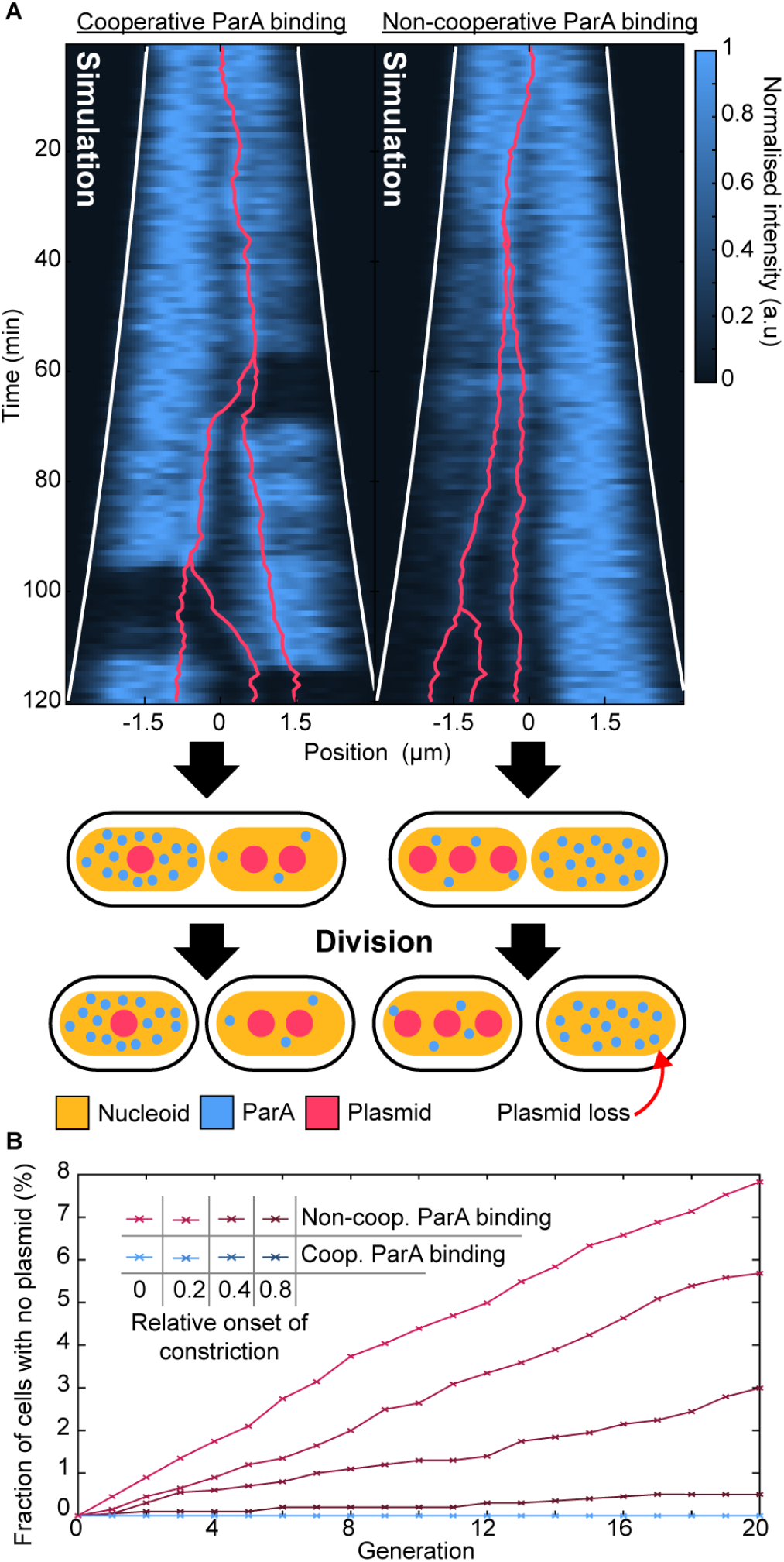
Cooperative ParA binding ensures plasmid stability. **(A) (**Top) Kymograph of a simulated cell with and without ParA cooperativity using the nucleoid profile shown in Figure 4E(i). (Bottom) Schematic representation of nucleoid, plasmids and ParA distribution before and after division for the two cases. In the non-cooperative scenario, only one cell inherits the plasmids. **(B)** ParABS system simulations were performed both with and without cooperative ParA binding across four distinct nucleoid constriction formation timings (0 birth, 1 division), spanning 20 generations. Each data point, representing ∼1,000 simulated cells, indicates the fraction of cells without plasmids in a particular condition and generation. At the end of each generation, the final plasmid positions are used to ascertain the copy number at birth distribution for the subsequent generation. Plasmid replication frequency, timing and the initial distribution of plasmids for the first generation were chosen from our experimental data. For all our simulations, we used the nucleoid signal of the cell from Figure 4E(i). This was then scaled temporally to place the onset of nucleoid constriction at the given relative cell ages. See SI Appendix for details.

To quantify the effect of this on plasmid stability within a population, we performed simulations of plasmid positioning over multiple generations. The initial plasmid copy number distribution was taken from our experimental data as were the timings of plasmid replication (see SI Appendix for details). We then measured the fraction of simulated cells with no plasmids at the end of each generation (Fig. 6B). Without cooperative ParA binding, we found detectable plasmid loss when the nucleoid begins to constrict early the cell cycle, with a loss rate per generation of 4 ·10^−3^ for the extreme case of constriction from birth (as we observed for rich media (Fig. 2G, S6F)). On the other hand, with cooperative binding, there were no incidences of plasmid loss across all 20,000 cell cycles simulated irrespective of the constriction timing, indicating a loss rate of at most ∼5 ·10^−5^ per generation, which is in the typical measured range (52). These findings support our hypothesis that ParA flipping is a mechanism to partition plasmids between the two lobes of the constricted nucleoid and is important for growth conditions in which nucleoid constriction begins early in the cell cycle.

## Discussion

Over the past several decades, extensive research on the ParABS system has shed light on numerous aspects of its function and role in DNA segregation, most notably the *in vitro* reconstitutions of the systems of the F and P1 plasmids (27, 28, 53). Even so, surprises remained as evidenced by the recent discovery that many ParB proteins are CTPases that act like DNA clamps, loading at the *parS* sites and subsequently sliding along the DNA (9–16). At the same time, techniques such as super resolution microscopy and chromatin immunoprecipitation (ChIP-Seq) have deepened our understanding of how ParA and ParB behave within cells (21, 25, 40, 44, 49, 52, 54). However, how the *in vivo* dynamics of ParA connect with cargo positioning has remained unclear. In particular, the flipping behaviour of ParA that has been observed for the plasmids F and TP228, and of *Vibrio cholerae* have not been explained (5, 34, 35). The current theory that oscillations arise from each plasmid following a self-created retracting gradient of ParA across the cell that fills in behind them (23, 26, 32, 33) is not consistent with our *in vivo* observation that ParA migration is demarcated by the mid-cell and is only correlated to plasmid mid-cell crossing events, with the other plasmids in the cell seemingly unaffected by the appearance or disappearance of ParA (Figs 2 & 3). We also discovered that the localisation of ParA to one cell half, and its flipping between cell halves, occurs more frequently as the nucleoid constricts (Fig. 2). Based on these observations, we propose that ParA binds to the nucleoid cooperatively and have shown here how this, along with constriction of nucleoid, can reproduce the *in vivo* behaviour. Note that while our simulations are not quantitative in the sense that there are unconstrained parameters, the fact that we can qualitatively reproduce all our measurements still provides strong support for the model.

Evidence of cooperative ParA binding to non-specific DNA has already been found for several systems including P1 plasmid, *V. cholerae* and *T. thermophilus* (5, 35, 36, 50, 51). Cooperative binding, albeit to the membrane, also occurs for the MinD, a relative of ParA, and is considered a key requirement for pattern formation in that system (55–57). Moreover, older *in vitro* studies showed that ParA, including from F plasmid, can form filaments (4, 29, 50, 58, 59). While these are not believed to occur *in vivo*, it nonetheless indicates the potential for self-interaction beyond dimer formation. Indeed, F-plasmid ParA was proposed to bind cooperatively to the promoter of its *par* operon through protein-protein interactions of its highly flexible winged-HTH domain (60). As for the molecular mechanism of the cooperativity, our model does not specify the details but we speculate that bound ParA-ATP dimers may promote the conformational change needed for cytosolic dimers to become competent for DNA binding (5, 6). Understanding this would likely have relevance across the diverse family of ParABS and related systems.

ParA asymmetry and flipping between nucleoid lobes occurs after the nucleoid begins to constrict. This led us to propose that flipping constitutes a second mode of action of the ParABS system that focuses on the *partitioning* of plasmids between nucleoid lobes as distinct from their *positioning* within each lobe and before nucleoid constriction. The flips establish a sharp gradient of ParA across the constriction from the nucleoid lobe with more plasmids towards the lobe with fewer. This gradient then directs the movement of a plasmid across the constriction, thereby acting to equalise the plasmid number in each lobe. Without such a mechanism, the difficulty of crossing the constriction would hinder or even prevent plasmid inheritance by the daughter cells. Our simulations supported this reasoning and showed a significantly higher plasmid loss rate in the absence of cooperative ParA binding, and therefore also ParA flipping.

In conclusion, this study has taken an important step toward unravelling the complexities of the ParABS system. While we have focused on F plasmid, our results are very likely also relevant for chromosomal ParABS systems, for which ParA flipping has also been observed (5, 61, 62). Our work also narrows the gap between ParA and its distant homologue, the cell division protein MinD, which also oscillates within the cell and binds cooperatively, but to the membrane. We expect that future studies will discover further commonalities within this diverse family of ATPases.

## Material and Methods

### Strains and growth condition

All experiments primarily use strain DLT3053 (44), a derivative of the *E. coli* K-12 strain DLT1215 (63) containing the HU-mCherry fusion to visualise the nucleoid. For the majority of our study, we use the miniF plasmid pJYB249 (miniF *parA-mVenus parB-mTurquoise2*) that carries functional fluorescent fusion to ParA and ParB under the native promoter (40). Figure S1G-I also presents data of the strain DLT1215 and plasmid pJYB234 (miniF *parB-mVenus*) from our earlier publications (64). These and similar fusions have previously been shown not to affect plasmid stability or expression levels (43, 44, 52, 65). The ParA dynamics we observe are also similar to that observed for a chromosomally-carried ParA fusion (34). Overnight cultures were grown at 37 °C in LB-Media containing 10 µg/ml thymine + 10 µg/ml chloramphenicol. For microfluidic experiments (see below) the following growth media were used: M9-Glycerol: (1x M9 salts + 0.5% glycerol + 0.2% casamino acids + 0.04 mg/mL thymine + 0.2 mg/mL leucine); M9-L-Alanine: (1x M9 salts + 0.2% L-alanine + 0.04 mg/mL thymine + 0.2 mg/mL leucine + mg/ml thiamine; LB: (LB + 0.04 mg/mL thymine).

### Microfluidics

Like the original mother machine (66), our design consists of a main channel through which nutrient media flows and narrow growth-channels in which cells are trapped. However, we follow (67) and include (i) a small opening at the end of each growth channel (ii) a waste channel connected to that opening to allow a continuous flow of nutrients through the growth channels (iii) an inverted growth-channel that is used to remove the background from fluorescence and phase contrast. We used a silicon wafer with this design to create the mother machine. We poured a polydimethylsiloxane (PDMS) mixture composed of a ratio of 1:7 (curing agent:base) over the wafer and let it rest at low pressure in a degasser for ∼30 min to remove air bubbles inside. The PDMS was then baked at 80 °C overnight (∼16 h). The cured PDMS was peeled off the wafer. Before imaging, the chip is bonded to a glass slide using a plasma generator (30 s at 75 W) and subsequently baked for a further 30 min at 80 °C, while the microscope is prepared.

### Microscopy

We used two Nikon Eclipse Ti-E microscopes with 100x oil-immersion objectives (configurations and settings in Table S2). All experiments were carried out using a mother machine except for those shown in Figure S4 and S7, which used agarose pads. For all experiments, overnight cultures grown on LB media were inoculated into the respective media used in the subsequent experiment 4 hours prior to the start of the experiment (2 hours for LB-condition) at 30 °C.

For mother machine experiments, cells were then loaded into the chip through the main channel and the chip was placed into a preheated microscope at 30 °C. The cells were constantly supplied with fresh media by pumping 2 μL/min of either of the three media that defined above, through the microfluidic chip. Cells were grown for an additional 2 hr inside the microscope before imaging began. Based on the specific (instantaneous) growth rate (Fig. S1F), this was sufficient time for cells to adapt to the device and the media. Cells were imaged at 1 minute intervals for approximately 72 hours (M9-Glycerol) and 24 hours (M9-L-Alanine & LB). Phase contrast, RFP-signal, CFP-signal and YFP-signal were captured for M9-Glycerol. CFP-signal was not captured for LB and M9-L-Alanine.

The short wavelength excitation of mTurquoise2 (CFP channel) can lead to cytotoxic effects (68). At the exposure used, we did not observe statistically significant incidences of cell cycle arrest or cell death (in any case such cells would be excluded by our segmentation pipeline). However, the mean cell cycle duration was 13% longer than what we previously observed while studying plasmid dynamics of F plasmid pJY243 (miniF, parB-mVenus) without CFP excitation (31, 64). Cells were also 26% longer at birth (Fig. S1G) and had a correspondingly greater plasmid copy number (Fig. S1H). However, growth appeared otherwise unperturbed and we observed no abnormal phenotype. Most importantly, plasmid positioning at the same copy number was consistent between the two datasets (Fig. S1I). We therefore believe that blue light excitation, which is difficult to avoid with the required three-color imaging, did not fundamentally affect our results.

For imaging on agarose pads (Figures S4 & S7), pads were prepared using M9-Glycerol media and 1% agarose. For the pads in Figure S7, either 200 µg/ml rifampicin or 200 µg/ml chloramphenicol was added and in the case of Thymine depletion, M9-Glycerol media, without the addition thymine, was used. 2 µl of culture was placed onto each pad prior to imaging. For the day cultures involving antibiotic pads, the same concentration of antibiotics was mixed into the culture right before placing it on the pad. In thymine depletion experiments, cells were first centrifuged to substitute thymine-containing M9-Glycerol media with the thymine-free version. Imaging commenced within 15 minutes after the cells were transferred onto the pad.

### Image processing

Cell segmentation was performed using a previously described image analysis pipeline (31). In addition, detected fluorescent foci were tracked using our recently developed ^★^ Track algorithm (https://gitlab.gwdg.de/murray-group/StarTrack, commit 38141270) (64). See SI Appendix for details.

### Kymographs and Normalisation

Single cell kymographs of an individual cell cycle (e.g. Fig. 1A) are stacked line profiles generated by averaging the signal across the cell’s short axis on each frame. Each kymograph shows the full cell cycle from birth to division unless stated otherwise. To ensure consistent visibility of the signal throughout the cell cycle, regardless of any change in the total signal, each line profile was normalised by dividing by its maximum value. Averaged kymographs combine line profiles across multiple cell cycles. Before combining, the line profiles were normalised in the same way. In Fig. 1D the averaged kymograph shows the average ParA signal before and after ParA flipping. Relative (from -1 to 1) rather than absolute position was used to allow the averaging of cells of different lengths (interpolation was done using the Matlab function *interp1*). In Fig. 2G the HU-mcherry signal from each cell cycle is averaged by normalising both the cell length and cell cycle duration. Each cell cycle starts at 0 (birth) and ends at 1 (division). Cells are normalised to length 1 at birth and length 2 at division and an exponentially adjusted length in between. This is an alternative to normalising cells to the same length irrespective of their relative cell age. For single-cell kymographs of simulated cells, the ParA location distribution is subjected to blurring in order to mimic the light diffraction inherent to microscopy. This was done using the Matlab function *smoothdata* using Gaussian smoothing with a window size of 10.

### Data, Materials, and Software Availability

Strains and plasmids used in this study are available upon request. Image data and Matlab scripts are available on Edmond repository at https://doi.org/10.17617/3.XOWNDX. The C++ code of the model is available at https://gitlab.gwdg.de/murray-group/hopping_and_relay/-/tree/ParA_Flipping.

## Acknowledgments

This work was funded through grant MU 4469/2 from the Deutsche Forschungsgemeinschaft (DFG).

R.K. received funding from the International Max Planck Research School for Environmental, Cellular and Molecular Microbiology. We thank Jean-Yves Bouet (Toulouse) for providing strains and plasmids.

