## Supporting Information for "Plasmid partitioning driven by collective migration of ParA between nucleoid lobes"

Seán M. Murray

**This PDF file includes:**

Supporting text
Figures S1 to S11
Table S1
SI References

### Supporting Information Text

#### Image processing

Our image processing pipeline for mother-machine experiments was described previously (1). It consists of four parts: (I) preprocessing, (II) segmentation, (III) cell and (IV) foci detection tracking. While Parts I, III and IV use custom Matlab scripts, Part II is based on SuperSegger (2), a Matlab-based package for segmenting and tracking bacteria within microcolonies (original code is available at https://github.com/wiggins-lab/SuperSegger), that we modified to better handle high-throughput data. SuperSegger employs pre-trained neural networks to segment cells by identifying their boundaries. It comes with a pre-trained model for E. coli which worked very well with our data. Therefore there was no need to train our own neural network. SuperSegger is capable of tracking cells however the tracking did not work properly with mother-machine images and so we developed our own method. Nevertheless, acknowledging that one of the main components of our pipeline, the segmentation, uses SuperSegger we refer to the entire pipeline as MotherSegger (code is available at https://gitlab.gwdg.de/murray-group/MotherSegger, commit cbc775b0).

In Part I, each frame of an acquired image stack is aligned (the offset between frames in x and y is removed). Afterwards the image stack is rotated so the growth channels are vertical. A mask of the mother machine layout is fitted to the phase contrast, using cross-correlation, to identify where the growth channels are located. Each growth channel is extracted from the image stack and the flipped inverted channel is subtracted to remove the background from both the fluorescence signal and phase contrast. The images are then segmented using Supersegger.

In Part III cells are tracked. Since cells cannot change their order inside the growth channel, they can be tracked by matching similar cell length between frames (starting from the bottom of the growth channels). To filter out potential segmentation errors, cell cycles that do not have exactly 1 parent and 2 daughters are excluded from analysis along with their immediate relatives (with the exception of those who are pushed out of the growth channel).

Part IV detects fluorescent foci within cells using the SpotFinderZ tool from Microbetracker (3), modified to our data structures. Tracks were inferred from the detected foci using ⋆Track (https://gitlab.gwdg.de/murray-group/StarTrack, commit 38141270) (4), which finds the optimal tracking accounting for false positive and negative detections. The used parameters were $D = 1.92 \cdot 10^{-4}$, $p_{neg} = 0.0822$, $p_{pos} = 0.0224$, $m = 3$, as found in that publication.

#### Detailed description of the computational Model

Our original model was described previously (1) and is an extension of the DNA-relay model (5) to include diffusion on the nucleoid (hopping) and basal hydrolysis of ParA and uses analytic expressions for the fluctuations rather than a second order approximation. Like the DNA relay model it is a 2D off-lattice stochastic model and updates positions in discrete time steps $dt$. The implementation was written in C++ and is available at https://gitlab.gwdg.de/murray-group/hopping_and_relay/-/tree/PaperParABS. It operates as follows. ParA associates to the DNA non-specifically in its ATP-dependent dimer state with the rate $k_a$ or through cooperative binding with the rate $k_r$. Once associated, ParA (i.e. ParA-ATP dimers) moves in two distinct ways: (i) Diffusive motion on the nucleoid with the diffusion coefficient $D_h$. This is an effective description of the movement of dimers due to transient unbinding events that allow them to ‘hop’ between DNA-strands. (ii) Between hopping events, each bound ParA dimer experiences the elastic fluctuations of the DNA strand it is bound to. This is implemented as elastic (spring-like) fluctuations around its current position. Dimers dissociate from the nucleoid due to either basal ATP hydrolyse at a rate $k_d$ or due to hydrolysis stimulated by ParB on the plasmid. The latter is modelled as a ParB-coated disc and ParB-ParA tethers form whenever the disk comes in contact with a ParA dimer. ParB-stimulated hydrolysis then breaks these tethers at a rate $k_h$, returning ParA to the cytosolic pool. The plasmid experiences the elastic force of every tethered ParA and moves according the its intrinsic diffusion coefficient $D_p$ and the resultant force of all tethers.

As in the DNA relay model, we only model three states of ParA: 'nucleoid associated' and 'cytosolic' and 'tethered'. Cytosolic ParA are assumed to be well mixed. This is justified based on the slow conformation changes needed to return it to a state competent for DNA-binding (6). No individual ParB molecules were modelled, rather the plasmid is treated as a disk coated with enough ParB that each nucleoid-bound ParA that makes contact with the plasmid instantaneously finds a ParB partner, therefore removing the need to model individual ParB.

The cell and nucleoid are modelled as a rectangle with dimensions $L \times W$. The positions of ParA and the plasmid(s), are updated as follows. Between hopping events, each nucleoid associated ParA dimer fluctuates about a home position $x_h$. The new position $x(t + dt)$ of each dimer is given by $x(t + dt) = x_h + \delta x$, where $\delta x$ is drawn with probability $p(\delta x, dt \mid x(t) - x_h)$ where $x$(t) is its original position and the normalised spring constant ($f/k_B T$ above) along each dimension is $1/\sigma_{x,y}^2$ and the diffusion coefficient $D_A$. During hopping events $x(t)$ and $x_h$ are both offset by a value drawn from a Gaussian distribution with $\mu = 0$ and $\sigma = \sqrt{2D_h dt}$ for both dimensions. The displacement of the plasmid is determined similar to each ParA dimer but according to the resultant force acting on it. This resultant force vector has an effective spring constant equal to the spring constant of a single tether times the number of tethers and acts towards an equilibrium position $x_p(t) + \sum_{tethers} (x_h - x(t))/n$, where $x_p(t)$ is the plasmid position and the sum is over all $(n)$ tethers. We ignore the effects of Torque. The intrinsic diffusion coefficient of the plasmid is $D_p$. If the plasmid has no tethers attached then it moves by normal diffusion, with displacements drawn from a Gaussian distribution with mean $\mu = 0$ and standard deviation $\sigma = \sqrt{2D_p dt}$. The $x$ and $y$ components of all positions are updated independently.

In this study, we extended this model to include cooperative ParA binding, cell growth, an upper limit of ParA-ParB tethers per plasmid, an accurate model of the nucleoid and its impact on ParA mobility. The code is found at https://gitlab.gwdg.de/murray-group/hopping_and_relay/-/tree/ParA_Flipping. Cooperative ParA binding is implemented as follows: each nucleoid bound ParA recruits, with rate $k_c = k_{c0} n_{ParA-cytosol}$, a cytosolic ParA to the same position on the nucleoid ($n_{ParA-cytosol}$ is the number of ParA in the cytosol at that moment). Growth was implemented by increasing the long axis of the cell at a rate of $k_g$ per simulation step and repositioning all particles (ParA & Plasmid) to preserve their relative positions within the cell.

To incorporate the nucleoid into our simulations, we take 2D HU-mCherry nucleoid signals from a cell cycle like the one shown in Figure 4E(i). The simulation is updated with the most recent nucleoid signal as the simulation progresses. Between updates, the signal is scaled with cell growth. The nucleoid signal acts as a potential on ParA dimers that are diffusing or hopping on the nucleoid. Thus, in regions where the nucleoid profile is flat, ParA dimers freely diffuse, whereas in the regions with a gradient, dimers are biased to move in the direction of more nucleoid signal, reflecting that they are more likely to hop to higher DNA-density regions. Note that the elastic fluctuations that dimers experience while they are bound is not affected. The displacement of the home position $x_h$ of each

ParA dimer when a nucleoid is present is implemented using a second-order approximation (7), as in the DNA relay model. The new home position of the dimer at time $t + dt$ is given by:

$$x_h(t + dt) = x_h(t) + R_0 + \frac{D_h}{k_B T} F dt + \frac{D_h^2}{2(k_B T)^2} FF'(dt)^2 + \frac{D_h}{k_B T} F' R_1$$

The initial three terms represent first-order approximations, while the final two terms are further expansion of the stochastic equations in (7). Here, $R_0$ and $R_1$ are random displacements following Gaussian distributions with $\mu = 0$ and $\sigma = \sqrt{2D_h dt}$ and $\mu = 0$ and $\sigma = \sqrt{(2/3)D_h(dt)^3}$ respectively, correlated as $< R_0 R_1 >= D_h(dt)^2$. The effective force $\frac{F}{k_B T}$ is calculated by multiplying the gradient (in 2D) of the nucleoid (linearly interpolated) at the current position $x_h(t)$ by the factor $p_n$, which defines the bias of ParA to move up the nucleoid gradient. Using the nucleoid signal of the cell cycle shown in Figure 4E(i), we conducted simulations varying $p_n$ and identified the value that resulted in a ParA distribution that most matched the actual ParA distribution observed *in vivo*. If the chosen value is too high, ParA accumulates excessively in regions with high nucleoid signal. If the chosen value is too low, the ParA distribution is too little affected by nucleoid density and no drop in the ParA signal is observed at the site of nucleoid constriction.

**Simulations for Plasmid Stability**

This section details how simulations were conducted to ascertain plasmid loss (Fig. 6B). Simulations were conducted across four unique nucleoid constriction formation timings (0 birth, 1 division) over a course of 20 generations. Each data point comprises data from at least 1000 simulated cell cycles, each lasting 120 minutes and starting with the average newborn cell length of 3 µm. The initial distribution of plasmids for generation 1 was derived from our experimental M9-Glycerol growth condition data. At the end of each simulated cell cycle, plasmid inheritance by the offspring was determined based on the number of plasmids in each cell half. This defined the distribution of plasmid numbers at birth for the next generation. This distribution is then multiplied by 1000 to determine the number of cells starting with a specific number of plasmids at birth (non-integer numbers are rounded up).

Plasmid replication was based on our experimental data. For every real cell cycle, plasmid replication timings in terms of relative cell age were determined. Each simulated cell was then assigned a random set of relative replication timings from the experimental pool, with the only condition being that the initial number of plasmids at birth matched. We found that plasmid replication is not random but occurs preferentially in the cell half with fewer plasmids. For copy number differences between cell halves of 1 and 2, 66% and 75% of replications respectively occur in the cell half with fewer plasmids. Differences of greater than 2 were too rare to quantify. We modeled this effect, and extended it to higher number differences $d$, by using a probability $p = \frac{d+1}{d+2}$ that replications occur in the cell half with fewer plasmids.

For all simulations, we used nucleoid signals from Figure 4E(i), for which constriction onset was identified at frame 24 out of 120. To emulate different constriction onset timings, we divided these nucleoid signals into two phases: pre-constriction onset (frames 1-23) and post-constriction onset (frames 24-120). These phases were then linearly scaled to different relative times of constriction onset. For instance, to simulate a relative onset of constriction of 0.8, the simulation uses the first 23 pre-constriction frames of the nucleoid signal, linearly distributed over the initial 80% of the simulated cell cycle and for the remaining 20%, the 97 post-constriction frames are used. The nucleoid signal is then used as described above to affect ParA mobility.

After all simulations were ran, the loss rate was calculated from the following function: $P_{loss} = 1 - \sqrt[20]{1 - f_{loss}}$ , where $f_{loss}$ is the fraction of cells of the last generation which did not contain any plasmids.

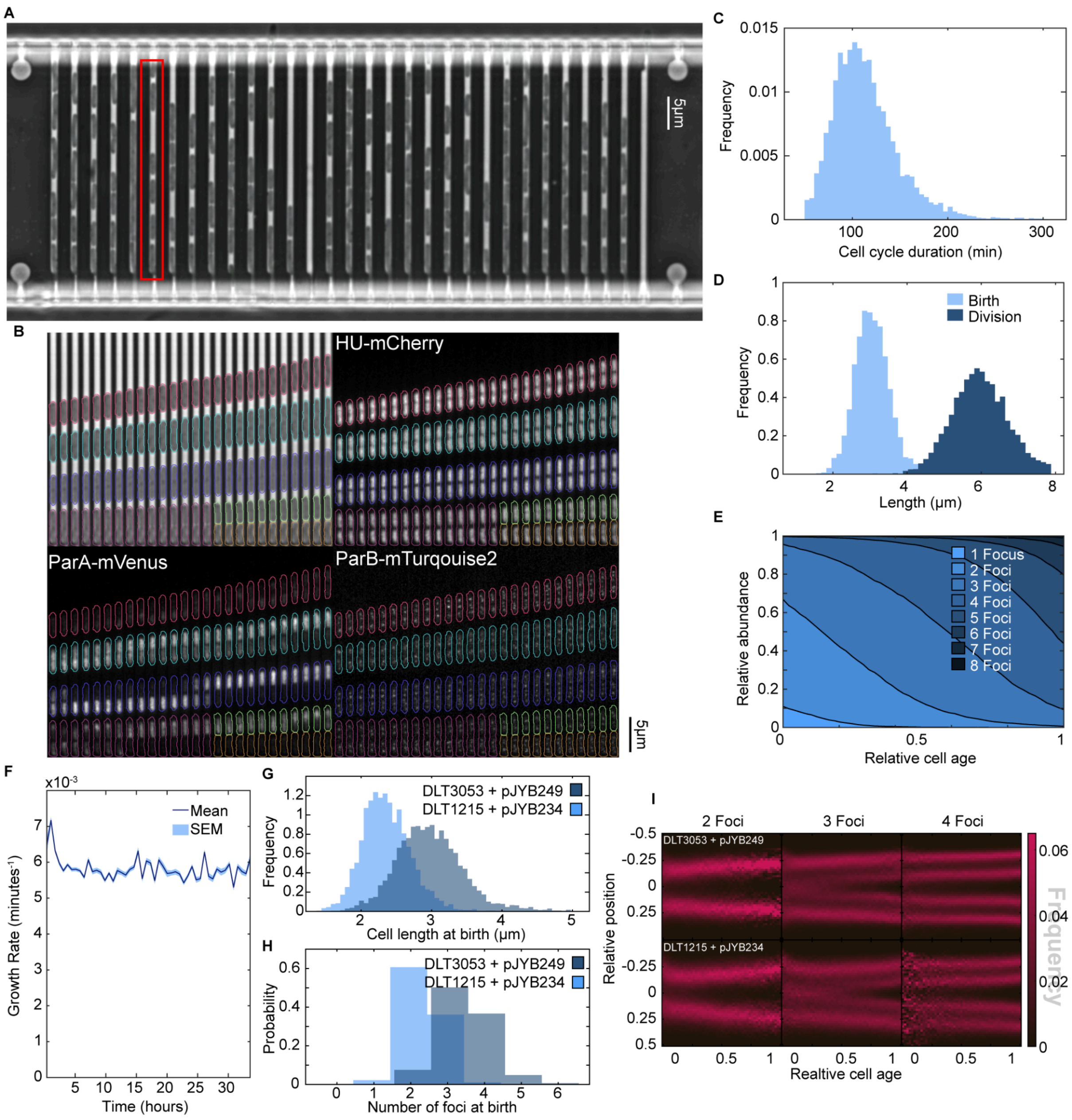


**Fig. S1.** Overview of a mother machine experiment. **(A)** Growth channels of one field of view. **(B)** Kymograph of one growth channel (highlighted in red in panel **(A)**) of the mother machine device. Phase contrast and 3 fluorescent channels are depicted with overlaid cell contours, colored by cell cycle. **(C)** Histogram of cell cycle durations (113.7 ± 34.8 min, mean ± SD). **(D)** Histogram of birth and division lengths (birth: 2.98 ± 0.49 µm, division: 5.98 ± 0.77 µm). **(E)** Relative abundance of ParB foci plotted against relative cell age. **(F)** Specific (instantaneous) growth rate of segmented cells plotted against imaging time. Note the deviation at the beginning and end is due to higher proportion of newborn and predivisional cells respectively and occurs irrespective of when the analysis begins. **(G)** Histograms of birth lengths from this study (DLT3053 + pJYB249; as in **(D)**) and from Köhler et al. 2022 (DLT1215 + pJYB234 (miniF *parB-mVenus*); mean ± std = 2.36 ± 0.34 µm, data from 4096 cell cycles) in which blue-light excitation was not used (see methods). The longer lengths of this study are due to the mildly cytotoxic effects of the short wavelength excitation (of ParB-Turquoise2). **(H)** Same as **(G)** but for the number of ParB foci at birth, showing a corresponding increase in copy number. **(I)** Average kymographs of the localization of 1 to 5 foci under both imaging conditions. The data from this study were generated from 5219 cell cycles.

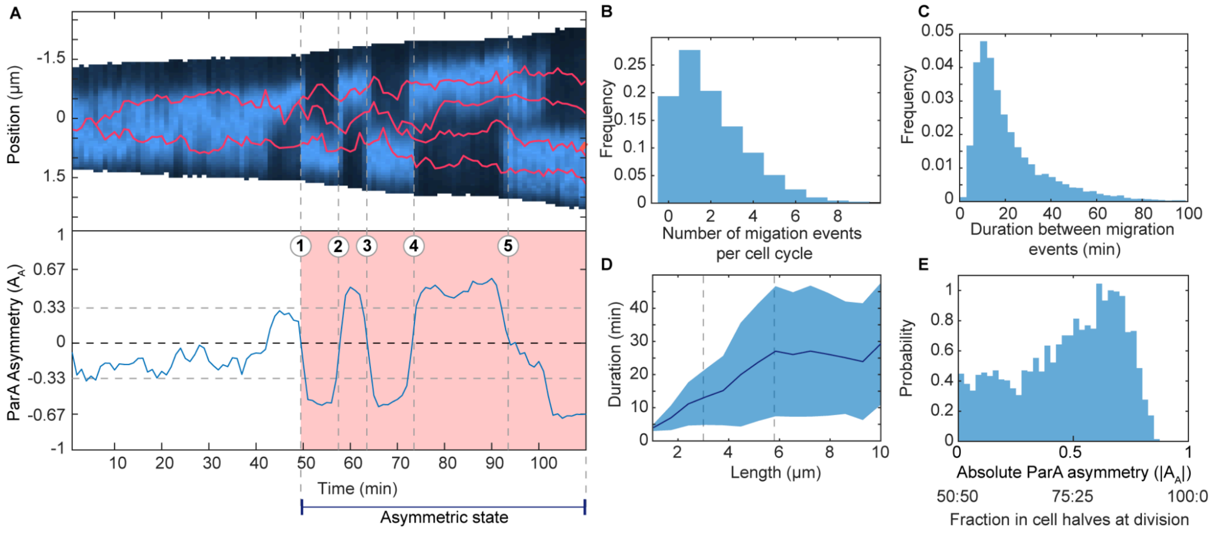


**Fig. S2.** Properties of ParA migration patterns. **(A)** Example of the annotation of migration events (flips) and when ParA is in an asymmetric state. Top: Kymograph of ParA-mVenus signal during the cell cycle of a cell. Red lines indicate plasmid trajectories obtained from tracking foci of ParB-mTurquoise2. Bottom: Fractional asymmetry of ParA. When the asymmetry exceeds the threshold indicated by the grey dashed lines (⅔ of the signal is localised in one cell half), the area between the zero-crossings is marked as ParA being in an asymmetric state (red shaded region). These zero-crossing points adjacent to asymmetric states (1)-(5) are defined as migration events of ParA (flips of ParA). **(B)** Distribution of the number of migration events per cell cycle. **(C)** Distribution of the duration between two flips. **(D)** The duration between migration events plotted against time since birth (mean ± SD). **(E)** The absolute asymmetry of ParA on the last frame from each cell cycle or the fraction of ParA in both cell halves directly before division. The panels **(B)** to **(E)** were generated from 5219 cell cycles.

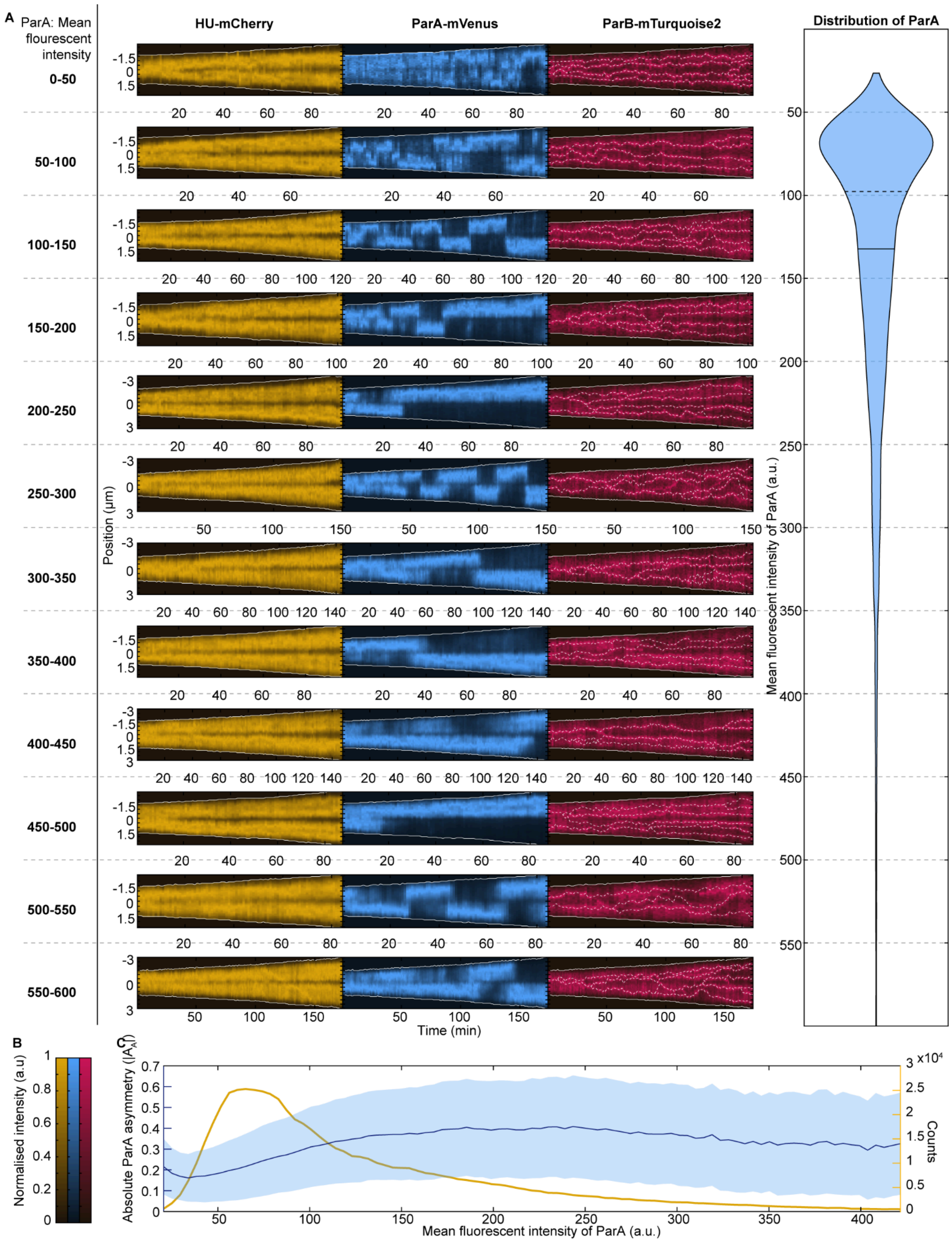


**Fig. S3.** Overview of the distribution of ParA within the population **(A)** ParA flips occur over the full range of ParA levels. Each example cell cycle shown has a mean ParA fluorescence intensity in the range indicated. Shown are kymographs of the nucleoid (HU-mCherry), ParA and ParB signals, Solid white lines indicate cell borders and dashed white lines show plasmid traces. Right: a violin plot showing the distribution of mean intensities across all cells. The horizontal dashed (solid) line indicates the median (mean) intensity. **(B)** Colour scales for **(A)**. **(C)** Absolute ParA asymmetry (mean ± standard deviation) plotted against mean ParA fluorescence intensity (blue line and shading) of each cell. The number of data points is shown in yellow. The violin plot in **(A)** and panel **(C)** were generated from 5219 cell cycles.

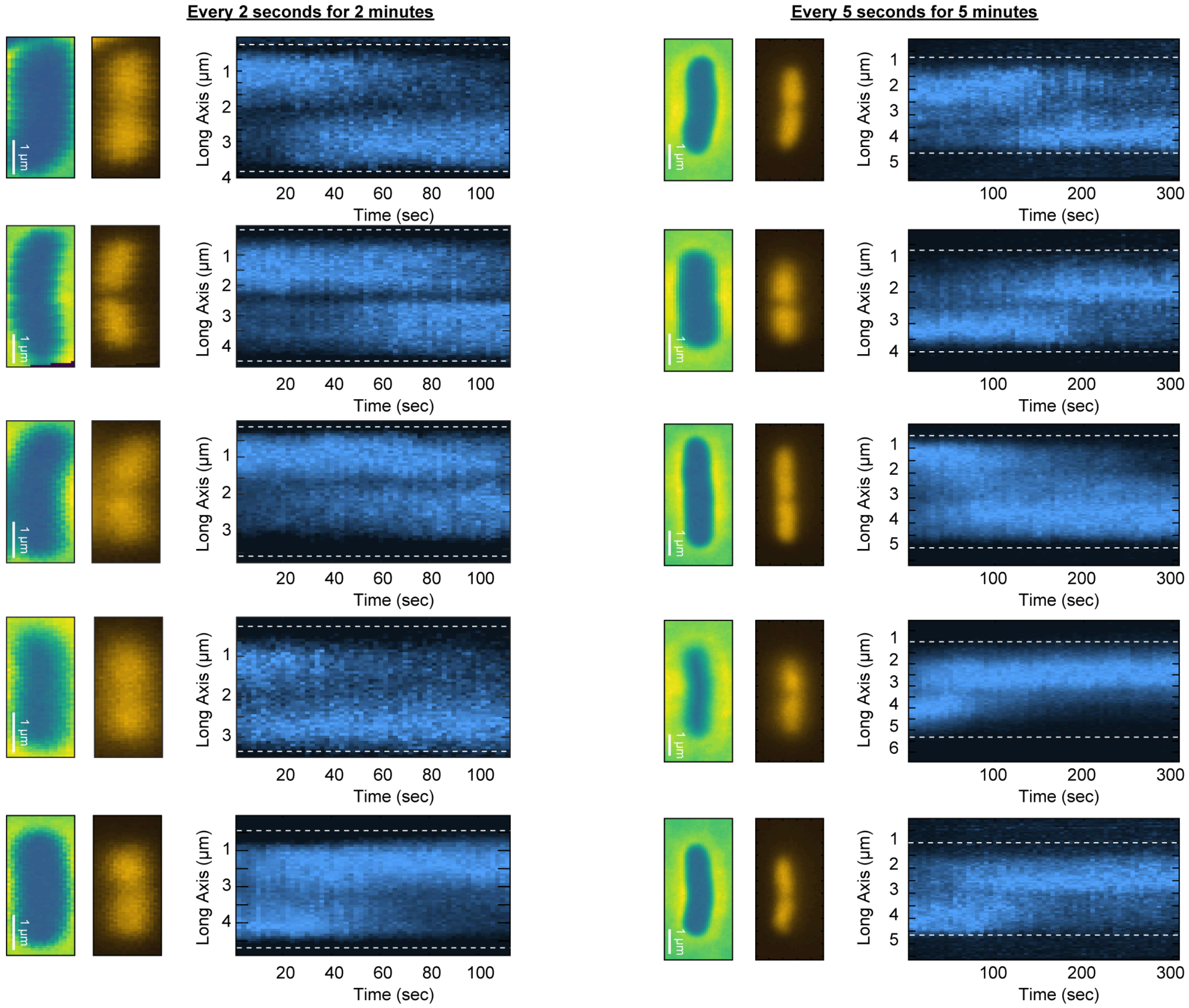


**Fig. S4.** ParA kymographs with high temporal resolution: Left: ParA signal was captured every 2 seconds over 2 minutes. Right: ParA signal was captured every 5 seconds over 5 minutes. Shown are, in order, phase contrast, nucleoid, and ParA profile along the long axis of the cell.

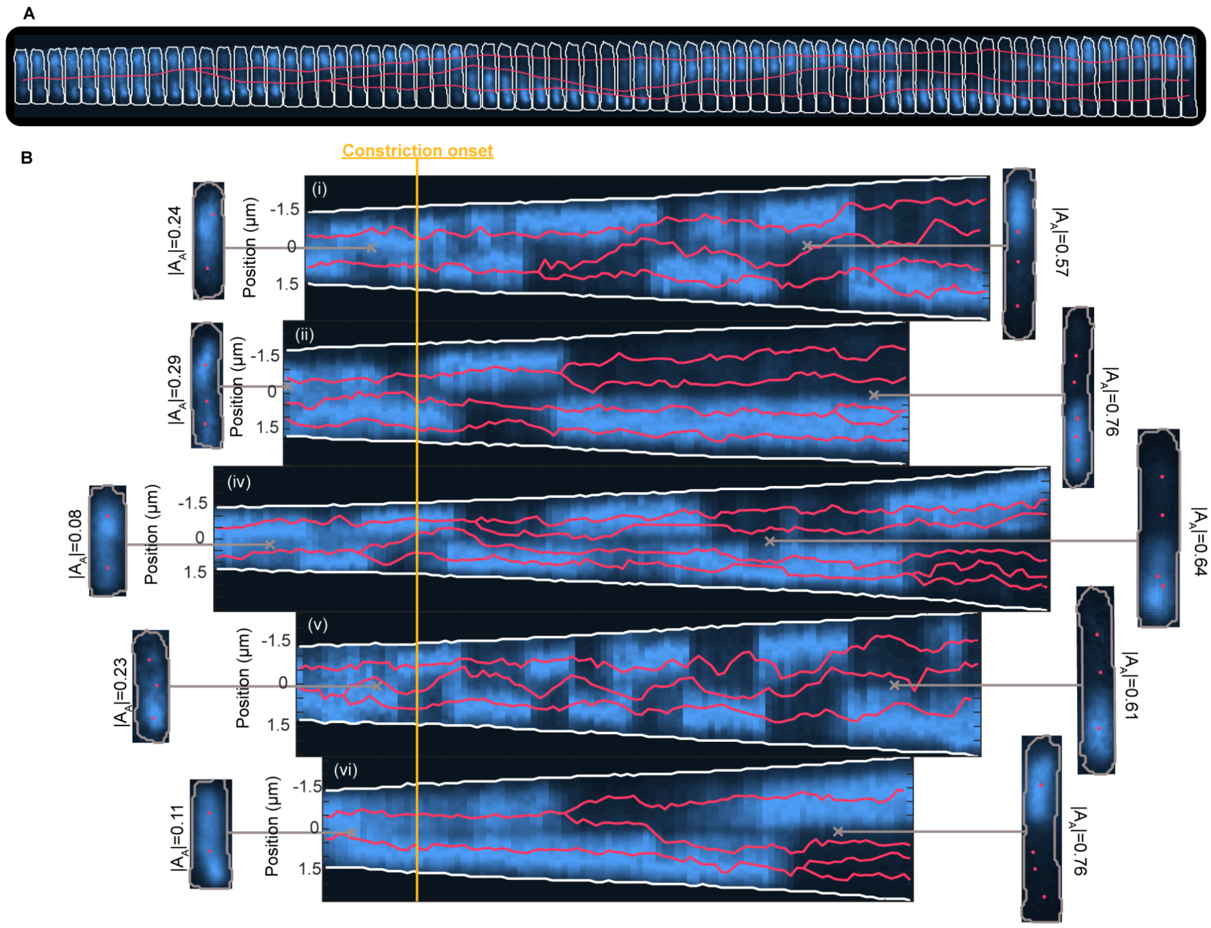


**Fig. S5.** Variability in ParA Dynamics. **(A)** Part of a cell cycle showing a plasmid apparently chasing a retracting ParA gradient (1 min intervals). This behaviour was rare, occurring in only approximately 3% of cells. **(B)** Presents kymographs of ParA from six cell cycles, synchronized at the point of constriction onset, indicated by the orange line. Red lines show the tracked movements of plasmids. Each cycle features single-frame overviews, each linked to its originating frame and annotated with the corresponding absolute ParA asymmetry ($|A_A|$).

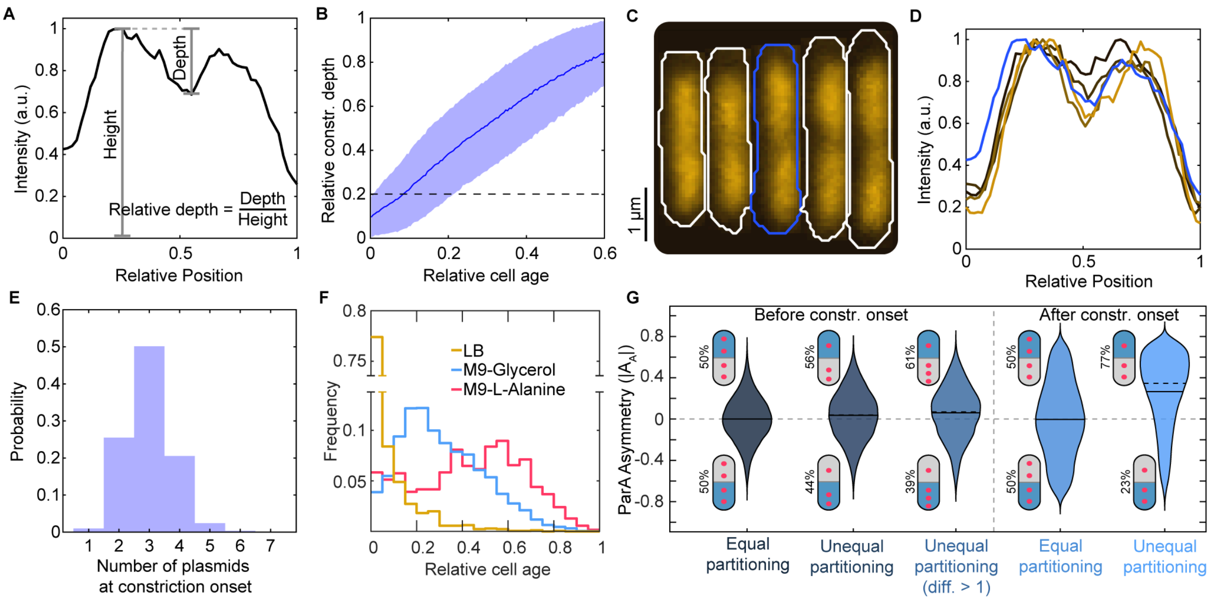


**Fig. S6.** Nucleoid constriction detection. **(A)** Example of an intensity profile of HU-mCherry (background subtracted). Highlighted are the height (max intensity) and the depth (distance between height and valley) and how to calculate the relative depth. **(B)** Average relative depth plotted against relative cell age; shade region is the standard deviation. The dashed line indicates the threshold depth for nucleoid constriction detection, defined as the 95% percentile of all relative depths on the first frame of each cell cycle (n=5219). This definition attempts to exclude random dips in the nucleoid and is based on the assumption that almost no cells have a genuine constriction at birth under slow growth conditions. **(C)** Example of nucleoid constriction formation. The cell outlined in blue indicates the initial instance where a nucleoid constriction was detected, and this feature persists in all subsequent cells. **(D)** Line profiles of the 5 cells shown **(C)**. The blue line corresponds to the cell with the blue outline. **(E)** Histogram of the number of plasmids at constriction onset. **(F)** Distribution of nucleoid constriction onset from the different growth conditions shown in Figure 2. **(G)** ParA asymmetry distribution before and after nucleoid constriction onset, binned by equal and unequal plasmid partitioning. For cells exhibiting unequal plasmid partitioning, cells are oriented such that positive ParA asymmetry corresponds to the majority of ParA being in the cell half with fewer plasmids as illustrated. Unequal partitioning is associated with significant ParA asymmetry after, but not before, constriction onset. Since the middle of an odd number of plasmids is typically positioned at mid-cell before constriction so that the plasmids are not truly unequally partitioned, we also show the ParA asymmetry of cells that have a disparity in the number of plasmids in each cell half of 2 or more. The distributions after constriction onset are also shown in Figure 3J.

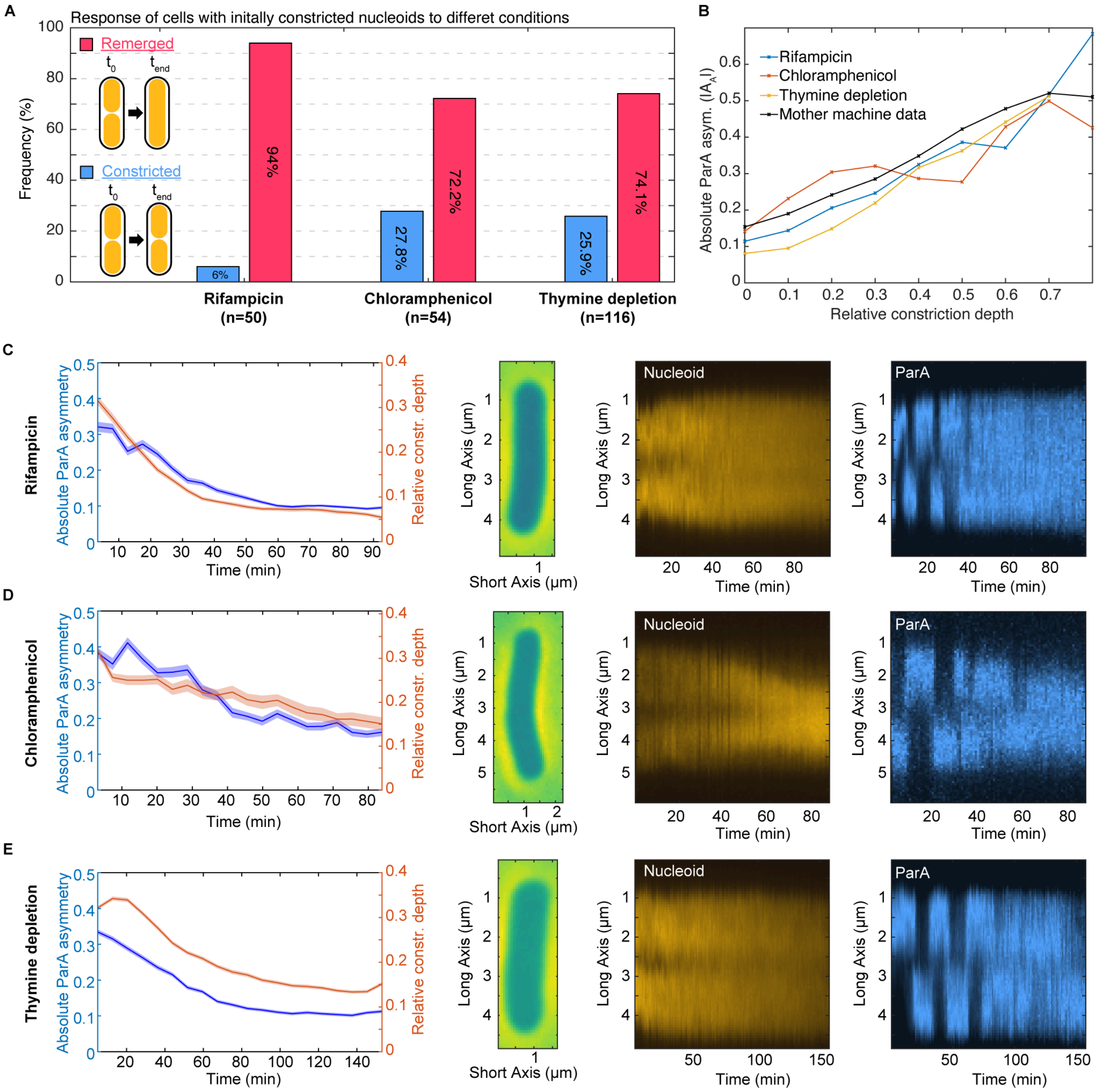


**Fig. S7.** ParA Dynamics in response to antibiotics and thymine depletion. **(A)** Bar chart showing nucleoid state (remerged or constricted) at the end of imaging for cells with an initially constricted nucleoid. Cells were exposed to Rifampicin (50 out of 112 cells initially constricted), Chloramphenicol (54/123 cells), and Thymine depletion (116/198 cells). Constriction is determined by the threshold described in Fig. S6. **(B)** ParA asymmetry versus relative constriction depth for initially constricted cells across the three conditions, compared with that of unperturbed cells as in Fig. 2F. **(C)** Left: Mean ± SEM of Absolute ParA asymmetry (blue) and relative constriction depth (orange) over time for initially constricted cells exposed to Rifampicin. Right: Phase contrast alongside nucleoid and ParA kymographs, displaying fluorescence from experiment start to end. **(D)** & **(E)** like **(C)** but for Chloramphenicol and Thymine depletion respectively.

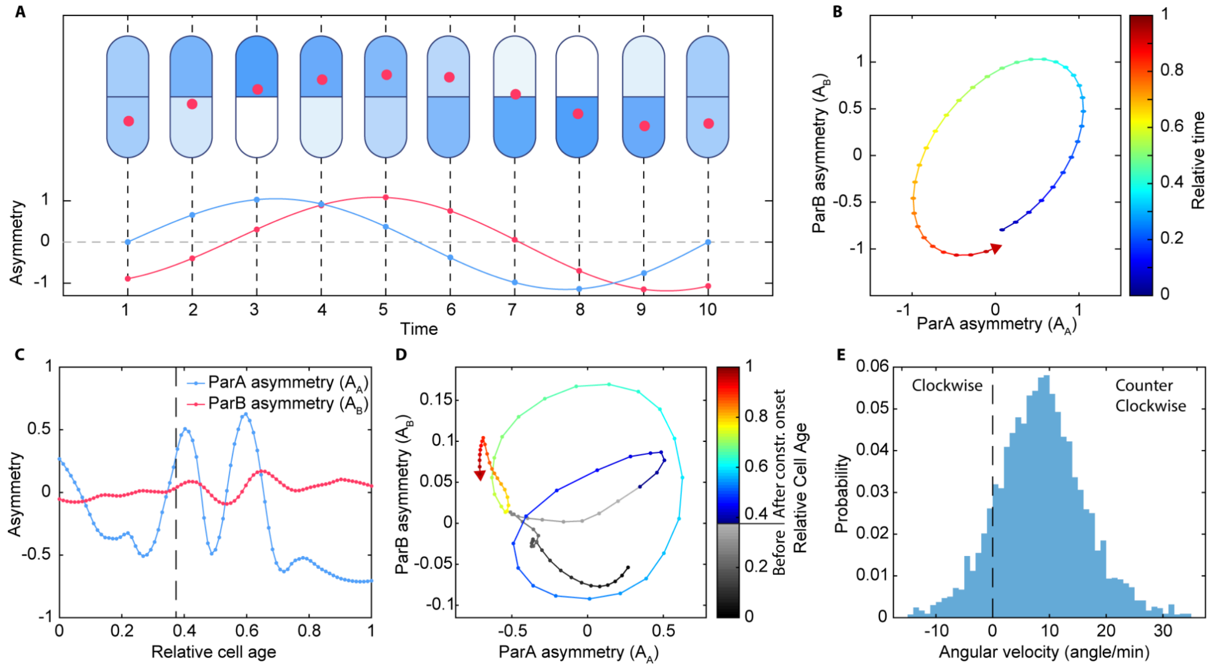


**Fig. S8.** The dynamics of ParA and ParB. **(A)** Top: A ParB-coated plasmid following ParA between cell halves. Bottom: Representation of the dynamics of ParA ($A_A$, blue) and ParB ($A_B$, red) asymmetry. **(B)** Both asymmetry values plotted against one another with color indicating time. The counterclockwise rotation indicates that ParB follows ParA rather than vice versa. **(C)** Asymmetry of ParA and ParB of a real cell cycle. The dashed line indicates the time point when onset of nucleoid constriction was detected. Both curves were smoothed using MATLAB's 'smooth' function, specifying 'rloess' as the method and 0.2 as the span. **(D)** The asymmetry of ParA and ParB plotted against each other creating a trajectory as the cell cycle progresses (same cell cycle as in **(C)**). The color of this trajectory shows relative cell age, with grey indicating data points prior to nucleoid constriction onset. **(E)** A histogram of the angular velocity after the nucleoid constriction was detected across all 5219 cell cycles. Angular velocities were determined from the trajectory of ParA and ParB asymmetry (example shown in **(D)**), by calculating the mean angle between all consecutive segments of the trajectory. A positive angle represents a counter-clockwise rotation, while a negative angle signifies a clockwise rotation between segments. The majority rotating counter-clockwise indicates that ParB (primarily bound to plasmids) follows ParA between the cell halves.

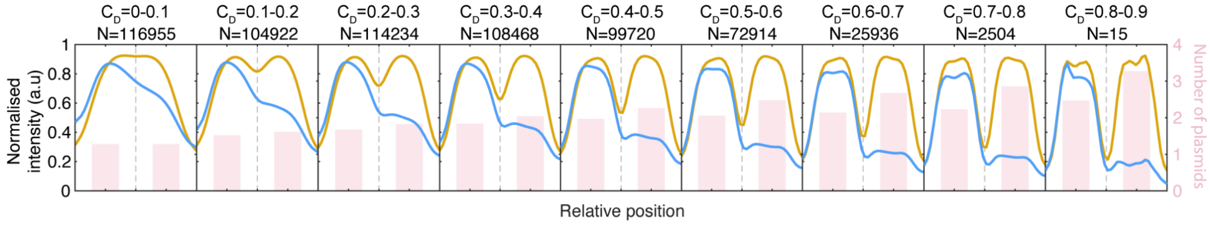


**Fig. S9.** ParA flips create a sharp gradient of ParA across the nucleoid constriction. Normalised profiles of nucleoid (orange) and ParA (blue, with max values set to 1 before averaging) for different constriction depths $C_D$ (0 indicates no constriction, 1 indicates complete separation/ no HU-mCherry signal between the lobes of the nucleoid). Cells were oriented such that the majority of ParA is in the same cell half. Light red bars in the background depict the average plasmid count in each cell half.

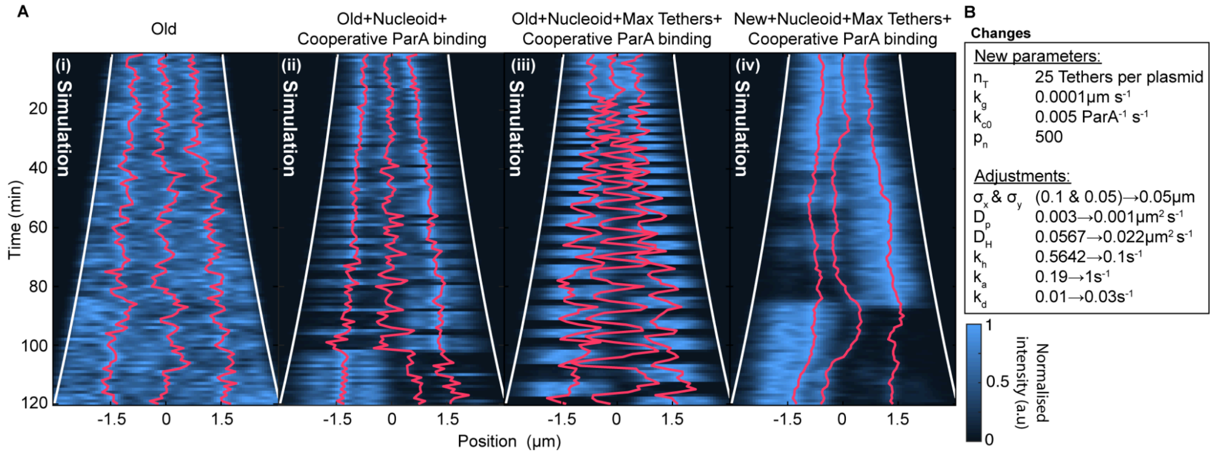


**Fig. S10.** New and adjusted parameters. **(A)** Kymographs of simulations with different parameters. (i): Parameters from Köhler et al., (ii): Parameters from Köhler et al. with the inclusion of a nucleoid and cooperative ParA binding, (iii): As in (ii) but with a limit on the number of ParA-ParB tethers, (iv) As in (iii) but with newly adjusted parameters. **(B)** New parameters that were introduced to the model: ($n_T$) Upper limit of ParA-ParB tethers per plasmid was set based on *in vivo* observations indicating no ParA foci at the plasmid's position. This also enhances plasmid mobility. ($k_g$) Growth rate of the cell. ($k_c$) Rate of cooperative ParA binding, ($p_n$) Attraction of ParA to the nucleoid was determined by matching the distribution of ParA within real and simulated cells. Adjusted parameters that were changed to mimic *in vivo* observations by reducing the mobility of the plasmid ($k_h$, $D_P$ & $\sigma_{x,y}$) and by reducing the frequency of ParA flips ($D_H$ & $k_d$) and by adjusting background levels of ParA ($k_a$).

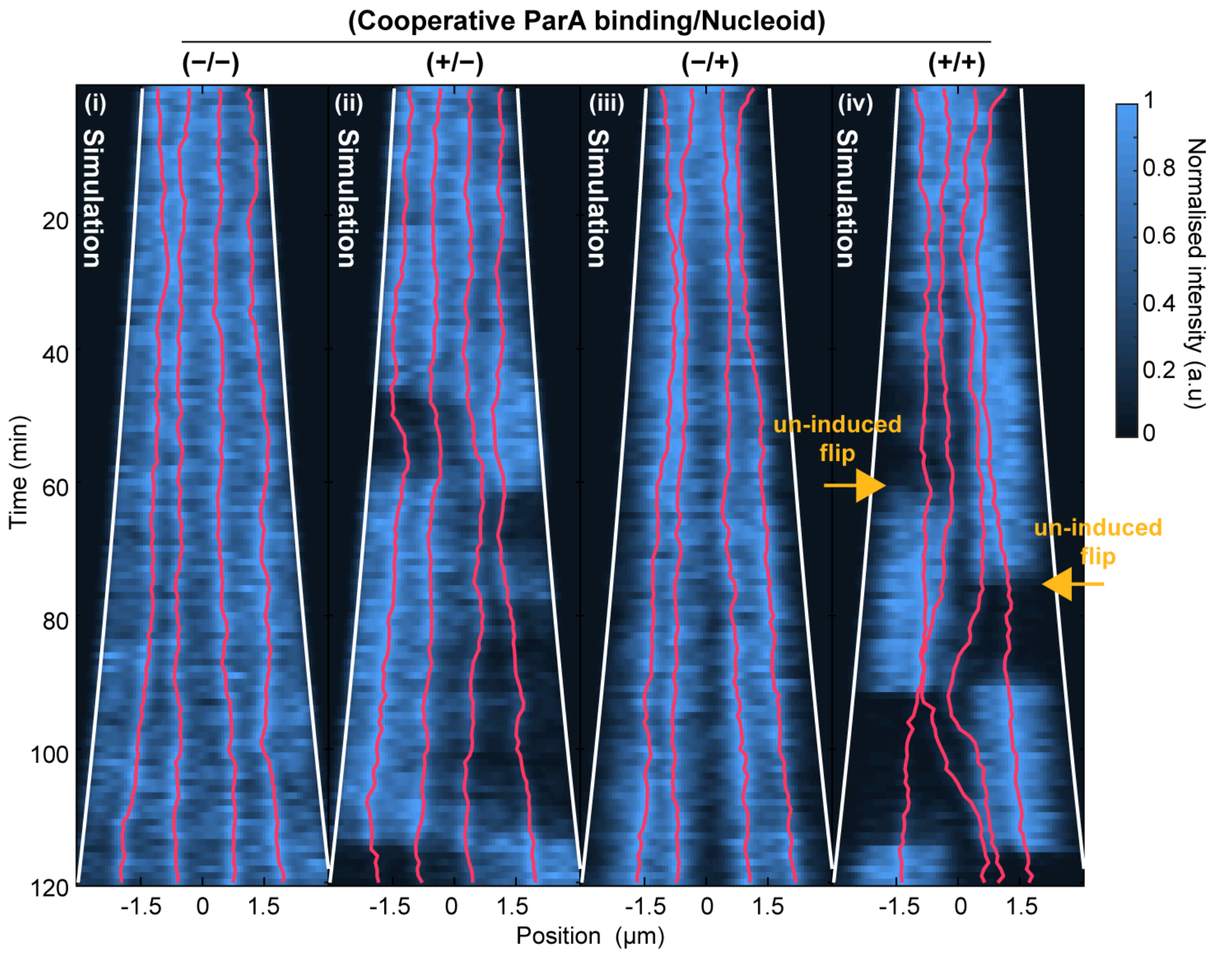


**Fig. S11.** Cooperative ParA binding causes instability of ParA at even plasmid numbers. Simulations with four conditions similar to Figure 4C but with four plasmids instead of 3. In (iv) the orange arrows indicate where the ParA does an un-induced flip (no plasmid crossing mid-cell preceding the flip).

**Table S1.** Microscopy settings.

Two Nikon Eclipse Ti-E microscopes with 100x phase contrast oil-immersion objectives (Plan Apochromat NA:1.45) were used with the given exposures and light intensities. Microscope #1 used an ORCA-Fusion C14440-20UP camera (Hamamatsu photonics) and a Lumen Dynamics XCT10A X-Cite fluorescence light source controlled by the software Nikon NIS Elements AR 5.20.01. Filters used for capturing the fluorescent signal are as follows (Exc,BS,Em): CFP (436/20, 455 LP, 480/40), YFP (510/20, 520 LP, 542/27), RFP (585/29, 605 LP, 647/57). Microscope #2 used an Andor Technology Zyla 4.2 CL10 sCMOS camera and Lumencor light engine Sola SE light source controlled by the Nikon NIS Elements AR 4.50.00. Filters used for capturing the fluorescent signal are as follows (Exc,BS,Em): CFP (438/24, 450 LP, 480/40), YFP (513/17, 520 LP, 542/27), RFP (575/25, 593 LP, 647/57).

| Figure | Intensity | Exposure | Frame rate | Microscope |
|---|---|---|---|---|
| 2 G,H | RFP:20%<br>YFP:20% | RFP:300ms<br>YFP:300ms | 1 minute | #1 |
| S1 G,H (DLT1215 + pJYB234) | YFP:20% | YFP:300ms | 1 minute | #1 |
| S7 A,B,D | RFP:30%<br>YFP:20%<br>CFP:20% | RFP:200ms<br>YFP:300ms<br>CFP:300ms | 1 minute | #2 |
| S4 left | RFP:50%<br>CFP:60%<br>YFP:50% | RFP:600ms<br>CFP:600ms<br>YFP:600ms | 2 seconds | #1 |
| S4 right | RFP:50%<br>CFP:60%<br>YFP:20% | RFP:600ms<br>CFP:600ms<br>YFP:600ms | 5 seconds | #1 |
| Rest | RFP:20%<br>CFP:50%<br>YFP:20% | RFP:300ms<br>CFP:500ms<br>YFP:300ms | 1 minute | #1 |
